# MAVS Safeguards Mitochondrial Integrity to Drive a Potent Intrinsic Antiviral Program

**DOI:** 10.1101/2025.04.28.651145

**Authors:** Vishal Sah, Poojitha Sai Potharaju, Karthika S Nair, Prangya Paramita Sahoo, Ravicanti Abhiram Pooja, Debasmita Basu, Dixit Tandel, K.S. Varadarajan, Santosh Chauhan, Anant Bahadur Patel, Krishnan Harinivas Harshan

## Abstract

Intrinsic antiviral defenses can restrict infection independent of paracrine interferon (IFN) signaling. While mitochondrial homeostasis is essential for immunity, its direct role in viral restriction remains incompletely understood. Here, we identify mitochondrial antiviral signaling protein (MAVS) as a central effector of a potent IFN-independent antiviral immunity orchestrated though mitochondrial safeguarding. We demonstrate that MAVS is critical for mitochondrial integrity as its loss leads to mitochondrial fragmentation, depolarization, and elevated mitophagy, alongside impaired bioenergetics and protein import. Mechanistically, MAVS maintains mitochondrial integrity by stabilizing the translocase of the outer membrane (TOM) complex and sustaining the abundance of its core components. Remarkably, this mitochondria-driven immunity restricts SARS-CoV-2 replication even under IFN-deficient conditions, and operates alongside the IFN pathways during JEV infection. Our findings redefine MAVS as a mitochondrial guardian that preserves organellar stability to enact host defense. These results highlight mitochondrial integrity as a fundamental determinant of broad antiviral immunity, integrating intrinsic and IFN-dependent mechanisms to counteract viral pathogenesis.

## INTRODUCTION

Interferon- and cytokine-mediated mechanisms dominate our understanding of the innate antiviral immunity. However, cells also deploy intrinsic, cell-autonomous antiviral defenses which are pre-existing, rapidly acting mechanisms that can restrict infection immediately within the infected cell, often before a paracrine interferon program is established(*1–3*). In classical usage, intrinsic immunity is mediated by restriction factors that directly inhibit discrete steps of the viral life cycle(*4, 5*), whereas “innate immunity” is often framed as a linear cascade of pathogen sensing – signaling – IFN/cytokine induction that coordinates local-to-systemic responses. How these two layers of antiviral defense intersect within infected cells remains incompletely understood. RNA sensing by cytosolic RIG-I–like receptors (RIG-I and MDA5) provides a useful lens through which to examine this intersection. Upon detection of viral RNA, these receptors signal through the mitochondrial antiviral signaling protein (MAVS), anchored on mitochondria and peroxisomes(*6, 7*). MAVS activation involves CARD-dependent oligomerization into higher-order assemblies that drive type I and type III interferon production and interferon-stimulated gene (ISG) expression(*8, 9*). Because this signaling output is essential for canonical innate immunity, diverse viruses antagonize MAVS or its assembly to blunt interferon responses(*10–14*). As MAVS-dependent antiviral signaling is primarily coordinated from mitochondria, this places the organelle itself at the center of host–virus interactions during infection.

Mitochondria are central hubs for innate antiviral defense, coordinating RNA sensing, signal transduction, organelle quality and metabolic adaptation during infection(*15, 16*). Intact mitochondrial homeostasis supports interferon (IFN)-dependent defenses. Many RNA viruses remodel mitochondrial architecture to create a permissive cellular state(*17–19*). Viral infection has been associated with mitochondrial fragmentation, fusion-biased elongation, loss of membrane potential, altered bioenergetics, and engagement of mitophagy(*20, 21*). These mitochondrial responses are highly dynamic and context-dependent, reflecting both virus-encoded activities and host attempts to maintain organellar homeostasis(*17, 19, 22–26*). Notably, many of these perturbations converge at the outer mitochondrial membrane and mitochondria–ER contact sites (MERCs/MAMs), where antiviral signaling complexes assemble and metabolic exchange is coordinated(*27, 28*). Despite extensive documentation of virus-induced mitochondrial remodeling, it remains unclear which host factors normally stabilize these mitochondrial territories during infection, how this buffering is lost, and whether preserving organellar integrity itself constitutes a barrier to viral replication.

One core function embedded within these same outer-membrane territories is mitochondrial protein import. It is a tightly regulated, multi-step process executed by translocases in the outer and inner membranes. Because mitochondrial fitness depends on continuous TOM/TIM-mediated import of nuclear-encoded proteins(23, 24), disruption of import is expected to destabilize respiratory function and membrane potential and to engage PINK1/Parkin-linked quality control, including mitophagy(25, 26). Of the many components of the TOM complex, the TOM receptors TOM20 and TOM70 form the precursor-recognition module at the outer membrane, a key interface between mitochondrial maintenance and antiviral signaling. Additionally, TOM70 has been reported to interact with MAVS during RNA virus infection and to facilitate recruitment of downstream TBK1/IRF3 signaling machinery to mitochondria(*29*). Consistent with its central role at the intersection of mitochondrial import and antiviral signaling, several RNA viruses directly target TOM70(*26, 30, 31*). MAVS signaling itself is coordinated from MERCs, placing antiviral signalosome assembly within outer-membrane territories that also govern protein handling and organelle quality control. Together, these observations place the TOM receptor module, and TOM70 in particular, at a nexus where import competence, mitochondrial integrity, and antiviral signaling intersect.

While studying the innate antiviral signaling during SARS-CoV-2 (CoV-2) infection, we identified a potent, IFN-independent intrinsic antiviral response that is derived from stable mitochondrial integrity. We further show that MAVS, best known for its role in interferon induction, has additional functions beyond canonical antiviral signaling. We present evidence to establish that MAVS is required to maintain mitochondrial network integrity, membrane potential, and oxidative phosphorylation during viral infections and otherwise. Its loss destabilizes core import machinery, remodels TOM organization, and elevates PINK1-linked mitophagy. Restoring MAVS re-establishes TOM receptor organization, import competence, and organellar fitness, reinstating antiviral resistance. Notably, this intrinsic immunity is capable of potently restricting viruses not just under conditions of IFN deficiency as in SARS-CoV-2 infections, but can also operate in parallel with canonical IFN pathways during infections by viruses such as JEV where IFN signaling is active. Together, our findings position MAVS as a mitochondrial guardian whose preservation of organelle integrity oversees an IFN-independent axis of antiviral defense, in addition to the canonical IFN-dependent programs.

## RESULTS

### MAVS is critical for maintaining mitochondrial structure and homeostasis

MAVS loss decapacitates the RLR signaling resulting in a muted IFN response upon viral infection. In MAVS KO Huh7 cells (generated using CRISPR/Cas9) we interestingly noticed that mitochondria were heavily fragmented in as compared against their WT-counterparts (puromycin selected post-vector transfection), which maintained normal mitochondrial population (Figures 1 A&B). Similar changes were noticed in multiple individual KO clones (Figures S1A) and also in primary mouse embryonic fibroblasts (MEFs) from Mavs^−^/^−^ mice (Figures S1 B&C), suggesting that this association of MAVS with mitochondrial morphology is well conserved across organisms. Given this significant morphological alteration, we next assessed mitochondrial membrane potential (Δψm). JC-1 staining revealed a significant reduction in Δψm in MAVS KO Huh7 and Mavs^−^/^−^ MEFs relative to their WT controls (Figures 1 C&D, Figures S1 D&E). Because ETC activity and Δψm are tightly coupled(*32*), we profiled mitochondrial bioenergetics using a Seahorse XF analyzer. MAVS-KO Huh7 cells showed significantly reduced basal/maximal OCR, collapsed spare capacity, and lower ATP-linked respiration (Figure S1F, Figures 1 E&F). Similar observations were made in primary Mavs^−^/^−^ MEFs as well (Figures S1 G&H). As a consequence of these changes, MAVS KO cells displayed markedly slower growth rates (Figure S1I). Failure to sustain efficient mitochondrial energetics in MAVS-KO cells prompted us to ask whether this bioenergetic collapse engages mitochondrial quality-control pathways. MitoGFP-RFP flux assay reported increased RFP-only (acidified) puncta per cell, indicating elevated mitophagy flux in MAVS KO Huh7 and Mavs^−^/^−^ MEFs (Figures 1 G&H, Figures S1 J&K). Elevated PINK1 phosphorylation and its colocalization with mitochondria (Figures 1I-L) indicated activated PINK1-dependent mitophagy in MAVS-deficient cells.

**Figure 1.**
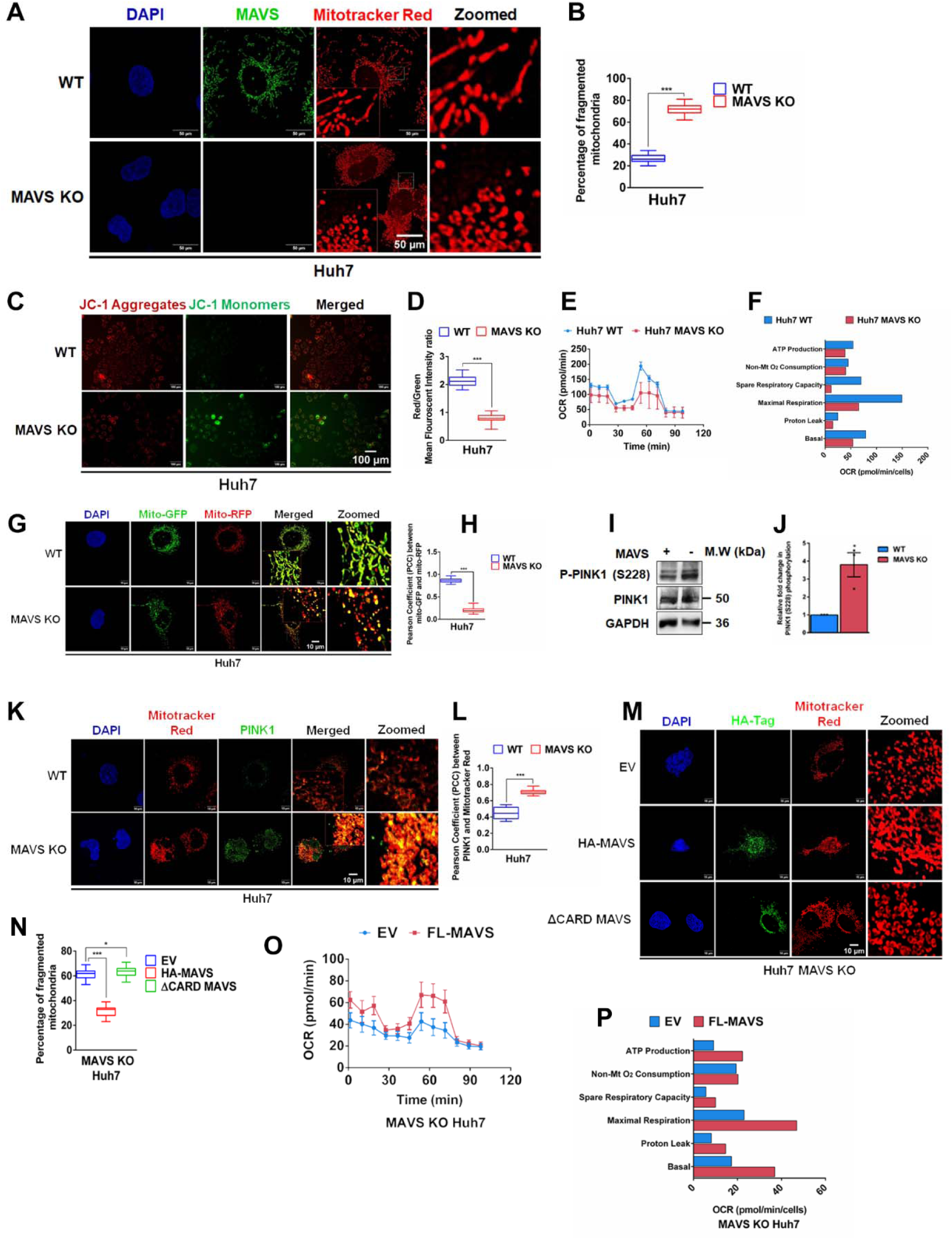
MAVS is critical for maintaining mitochondrial structure and homeostasis. **(A)** Confocal images showing mitochondrial fragmentation in Huh7 WT and MAVS KO cells, and **(B)** its corresponding quantification. Scale bars, 50 µm. **(C)** Epifluorescence microscopy images analyzing mitochondrial membrane potential in WT and MAVS KO Huh7 cells. Scale bars = 100 µm, and **(D)** quantitative representation of mitochondrial membrane potential from images in (C) (n = 100 cells). **(E)** Oxygen consumption rate (OCR) and **(F)** mitochondrial respiration parameters in WT and MAVS KO Huh7 cells measured by Mito Stress Assay and analyzed using the Seahorse XFe24 analyzer. **(G)** Confocal images showing mitophagy in Huh7 WT and MAVS KO cells transfected with a Mito-GFP-RFP reporter, and **(H)** corresponding quantification of mitophagy. **(I)** Immunoblot analysis of PINK1 phosphorylation in WT and MAVS KO cells, and **(J)** densitometric quantification. **(K)** Confocal images showing co-localization of PINK1 (green) with mitochondrial stained with Mitotracker Red (red) in WT and MAVS KO cells, and **(L)** corresponding quantification of co-localization. **(M)** Confocal images showing mitochondrial fragmentation in MAVS KO Huh7 cells transfected with FL-MAVS or ΔCARD MAVS, and **(N)** corresponding quantification. **(O)** OCR and **(P)** mitochondrial respiration parameters in MAVS KO cells transfected with FL-MAVS. Scale bars, 10 µm and n = 50 cells unless otherwise indicated. Data are presented as mean ± SEM, with *, ** and *** denoting p <0.05, 0.01 and < 0.001.

To confirm the association of MAVS with these changes, we re-expressed HA-MAVS in MAVS-KO Huh7 cells and reassessed mitochondrial integrity. As a signaling-defective control, we introduced a CARD-deleted mutant (ΔCARD-MAVS) that cannot oligomerize on the outer mitochondrial membrane. Full-length MAVS, but not ΔCARD mutant restored tubular mitochondrial networks (Figures 1 M&N) and mitochondrial bioenergetics (Figures 1 O&P). In MAVS competent cells, full-length MAVS supplementation resulted in enhanced mitochondrial network and bioenergetics (Figures S1 L-O). These data show that MAVS is necessary to maintain mitochondrial homeostasis.

### MAVS safeguards mitochondria by stabilizing TOM complex and protein import

The magnitude of mitochondrial defects upon MAVS perturbation suggested broad remodelling of mitochondrial programs. To define mitochondrial programs responsive to MAVS, we carried out deep RNA sequencing in MAVS-overexpressing Huh7 cells and in primary Mavs−/− MEFs. Using these perturbations across different animal models we tested whether shared mitochondrial pathways were consistently impacted. A mitochondrial enrichment analysis showed a strong correlation between MAVS and mitochondrial protein import. Several components of the translocase of the outer mitochondrial complex (TOM) were downregulated in MAVS KO MEFs, and upregulated in MAVS overexpressing Huh7 cells (Figures 2A & S2A). Their positive correlation with MAVS expression strongly suggested that MAVS is critical for the TOM complex assembly. These observations were further confirmed by immunoblotting (Figures 2 B&C). TOM20, and TOM70, two important accessory components of the TOM complex, were significantly underexpressed in MAVS KO cells, indicating selective loss of core import components. Additionally, TIM23, a key component of the translocase of the inner mitochondrial TIM23 complex, also showed compromised expression in these cells. Blue Native PAGE analysis showed reduced assemblies of TOM core and accessory components (Figures 2D&E). TIM23-containing assemblies were similarly reduced/redistributed, indicating compromised TIM23 complex integrity. Because TOM70 has been reported to interact with MAVS(*29*) and TOM20 associates with TOM70 within the TOM receptor module(*33*), we asked whether MAVS loss disrupts TOM20/TOM70 organization on the mitochondrial surface. High-resolution immunostaining with object-based quantification showed reduced TOM20 and TOM70 puncta densities (∼40% each) after normalization to mitochondrial area (Figures S2B–D). Object-based colocalization further revealed ∼50% fewer TOM20-TOM70 intersections and increased median centroid-to-centroid distances (Figures S2E–G), consistent with reduced TOM receptor clustering in the absence of MAVS. Since a major disorganization of TOM complex is likely to impact mitochondrial protein import, we assessed it directly using an MTS-GFP reporter. MAVS-KO Huh7 cells displayed increased accumulation of cytosolic GFP, comparable to treatment of WT cells with CCCP (a common mitotoxin)- (Figures 2F&G), consistent with impaired mitochondrial import. This import defect was rescued by MAVS re-expression: HA-MAVS restored mitochondrial targeting of the MTS-GFP reporter and reduced cytosolic GFP accumulation in MAVS-KO Huh7 cells, whereas the ΔCARD mutant failed to restore import (Figures 2H&I). This import defect is consistent with, and likely contributes to – the PINK1 accumulation and elevated mitophagy observed in MAVS-KO cells, given that PINK1 stabilizes on dysfunctional, import-incompetent mitochondria. Together, these data show that MAVS preserves TOM receptor abundance and organization to sustain mitochondrial import competence.

**Figure 2.**
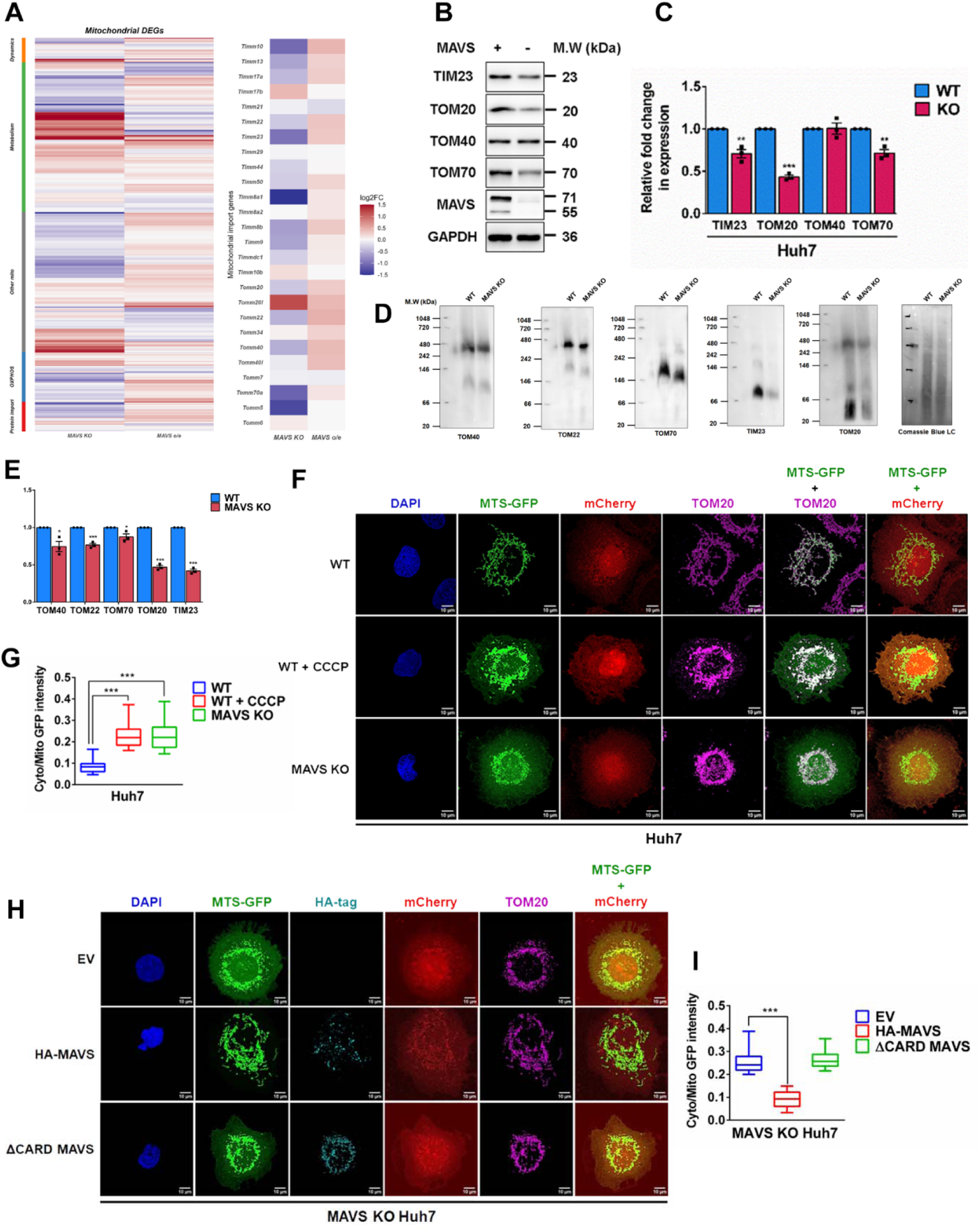
MAVS safeguards mitochondria by stabilizing TOM complex and protein import. **(A)** Heatmap showing the most significantly regulated mitochondrial DEGs, and DEG subset of genes responsible for mitochondrial protein import, between MAVS-supplemented Huh7 cells and MAVS KO MEFs generated via homologue mapping. Expression values are represented as Log_2_ Fold Change. **(B)** Immunoblot analysis of TIM23, TOM20, TOM40, TOM70, and MAVS levels in Huh7 WT and MAVS KO cells, and **(C)** the corresponding quantification of their relative levels. **(D)** BN-PAGE analysis of TOM40, TOM22, TOM70, TIM23 and TOM20 abundance in enriched mitochondria prepared from Huh7 WT and MAVS KO cells, and **(E)** corresponding quantification. **(F)** Confocal images showing cytosolic accumulation of MTS-GFP in Huh7 WT and MAVS KO cells, and **(G)** corresponding quantification of cyto/mito ratio. CCCP treatment was used as a positive control. **(H)** Confocal images showing cytosolic accumulation of GFP in Huh7 MAVS KO cells supplemented with FL-MAVS and ΔCARD MAVS, and **(I)** corresponding quantification of cyto/mito ratio. Scale bars, 10 µm and n = 50 cells unless otherwise indicated. Data are presented as mean ± SEM, with *, ** and *** denoting p <0.05, 0.01 and < 0.001.

### Viruses target MAVS aggregation to destabilize mitochondrial homeostasis

RNA viruses are known to target MAVS to blunt innate immune signaling(*11, 13, 34, 35*). Given that MAVS loss resulted in mitochondrial destabilization, we hypothesized that loss in MAVS oligomerization is associated with mitochondrial defects that mirror MAVS deficiency. Consistent with prior reports(*36, 37*) and our previous work(*38*), JEV infection robustly activated RLR–IRF3 signaling by 24 h post infection (Figure S3A), confirming engagement of the MAVS pathway under these conditions. We and others have previously reported that SARS-CoV-2 dampens IRF3 activation and IFN signaling(*10, 39–41*) We did not notice any change in MAVS abundance (Figures 3A&B; Figures S3B&C), but the infection failed to induce a type I interferon response (Figure 3C, Figure S3D)(*10*). We therefore tested whether SARS-CoV-2 perturbs MAVS oligomerization.

**Figure 3.**
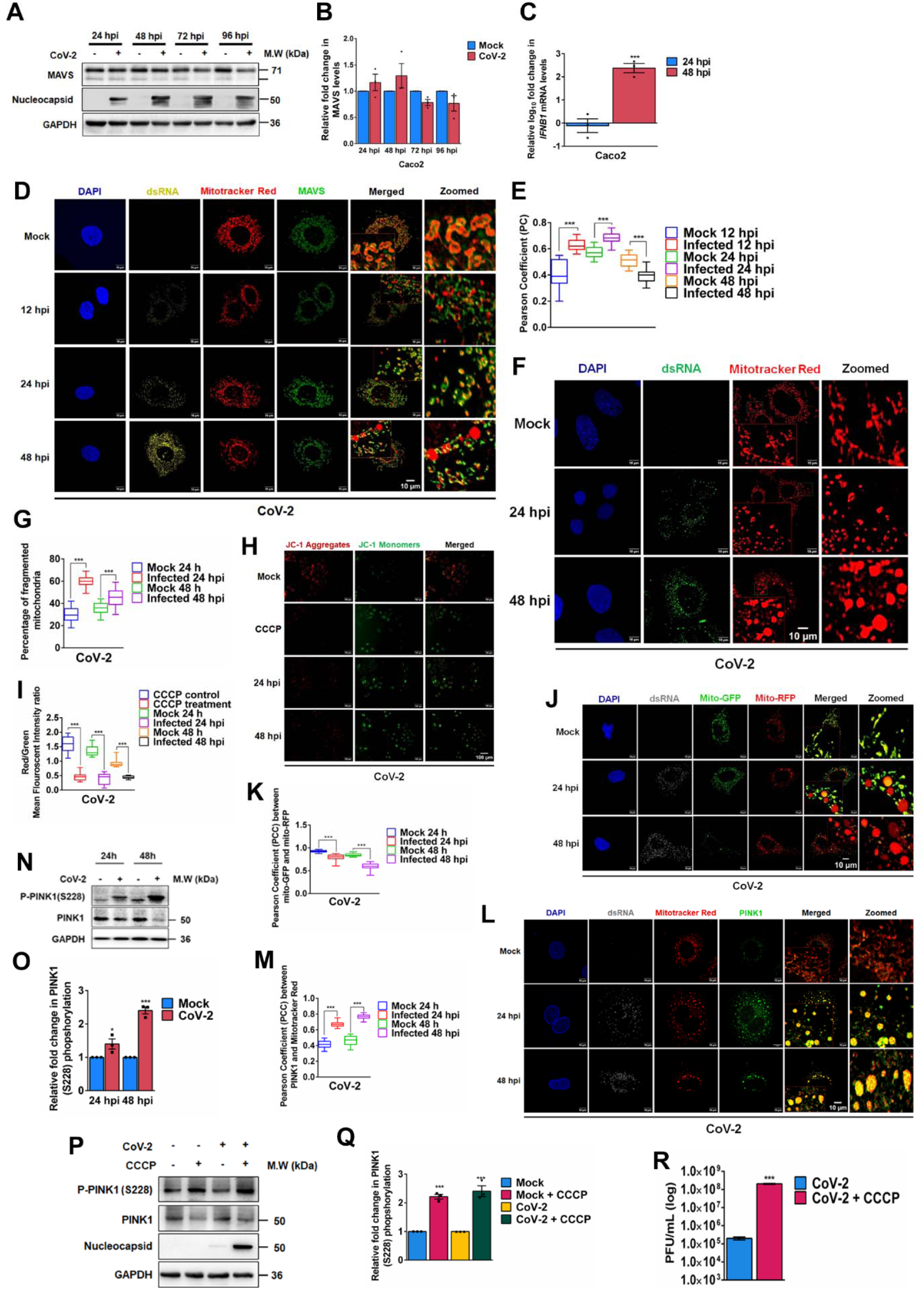
Viruses target MAVS aggregation to destabilize mitochondrial homeostasis. **(A)** Immunoblot analysis of MAVS and SARS-CoV-2 nucleocapsid protein expression levels shown at different time points upon SARS-CoV-2 infection in Caco2 cells and, **(B)** its densitometric analysis **(C)** qRT-PCR analysis showing relative fold change in *IFNB1* levels in Caco2 cells at 24- and 48-hours post-infection (hpi). **(D)** Confocal images showing MAVS aggregation in Huh7 cells infected with SARS-CoV-2, stained with Mitotracker Red and immunostained for dsRNA and MAVS. **(E)** Quantification of Pearson’s correlation coefficient (PCC) for MAVS-mitochondria co-localization from (D) (n = 250 ROIs). **(F)** Confocal images showing mitochondrial fragmentation in SARS-CoV-2 infected Huh7 cells. **(G)** Quantification of mitochondrial fragmentation from **(F)**. **(H)** Epifluorescence microscopy images analyzing mitochondrial membrane potential through JC1 staining at different time points post-SARS-CoV-2 infection (MOI = 1), with CCCP as a positive control. **(I)** Quantification of mitochondrial membrane potential from (H) (n = 100 cells). Scale bars, 100 µm. **(J)** Confocal images showing mitophagy in SARS-CoV-2 infected Huh7 cells transfected with a Mito-GFP-RFP reporter, and **(K)** corresponding quantification of mitophagy. **(L)** Confocal images showing PINK1 (green) co-localization with MitoTracker Red (red) in infected cells, and **(M)** quantification of PINK1 localization. **(N)** Immunoblot analyzing PINK1 phosphorylation (S228) in SARS-CoV-2 infected Huh7 cells, and **(O)** densitometric quantification. **(P)** Immunoblot analyzing PINK1 phosphorylation following CCCP treatment in uninfected or SARS-CoV-2 infected Huh7 cells. **(Q)** Densitometric analysis of PINK1 phosphorylation from (P). **(R)** Infectious viral titers from corresponding supernatants. Scale bars, 10 µm and n = 50 cells unless otherwise indicated. Data are presented as mean ± SEM, with *, ** and *** denoting p <0.05, 0.01 and < 0.001.

MAVS aggregation was quantified using a confocal colocalization-based assay previously described(*42*), with aggregation scored by the Pearson coefficient between MAVS and mitochondrial signals (Figure S3E). Poly(I:C) stimulation robustly increased MAVS aggregation, whereas the non-oligomerizing ΔCARD-MAVS mutant served as a negative control (Figures S3F&G) (*13, 43, 44*). SARS-CoV-2 infection failed to increase MAVS aggregation across the infection time course (Figures 3D&E). Notably, mitochondria in dsRNA-positive SARS-CoV-2 infected cells were largely devoid of MAVS puncta, indicating exclusion of MAVS from the mitochondrial network during infection (Figure S3H). We next asked whether this redistribution of MAVS during SARS-CoV-2 infection is accompanied by changes in mitochondrial organization. SARS-CoV-2 infection induced extensive mitochondrial fragmentation (Figures 3F&G), loss of mitochondrial membrane potential (Figures 3H&I), and increased mitophagosome formation (Figures 3J&K), accompanied by increased PINK1 accumulation and phosphorylation (Figures 3L-O). JEV and DENV have previously been shown to target MAVS and its aggregation(*45, 46*). Similar to SARS-CoV-2, JEV disrupted mitochondrial network integrity and homeostasis (Figures S3I-N), whereas DENV promoted mitochondrial elongation while largely preserving mitochondrial homeostasis, consistent with previous reports(*23, 24, 47*). These differences suggested a pro-viral role for mitophagy, prompting us to investigate this association. Interestingly, CCCP-induced mitophagy significantly enhanced SARS-CoV-2 replication (Figures 3P-R, Figures S3O&P), and JEV showed a similar increase (Figures S3Q&R). In contrast, CCCP treatment reduced DENV infection (Figures S3S&T), indicating that mitophagy promotes SARS-CoV-2 and JEV infection but is not beneficial – and may be detrimental to DENV under these conditions. These results indicated that targeting MAVS aggregation is means adopted by several viruses in their pursuit of subverting mitochondrial homeostasis. It is likely that DENV coordinates unique mechanisms to counter the mitophagy induction resulting from MAVS disaggregation.

### MAVS implements mitochondrial homeostasis to restrict RNA virus replication

Given that MAVS is a well-established antiviral adaptor, our finding of a mitoprotective role of MAVS suggested that MAVS is the link between the mitochondrial homeostasis and antiviral immunity. We tested whether boosting MAVS can restore mitochondrial integrity during infection and thereby limit viral replication by observing MAVS gain- and loss-of-function in parallel across infection models. Full-length MAVS supplementation, but not the ΔCARD mutant, significantly reduced SARS-CoV-2 replication across multiple cell lines and experimental systems (Figures 4A-D; Figures S4A&B). Consistent with a MAVS-dependent restriction, MAVS-KO cells supported markedly higher SARS-CoV-2 titers than WT controls (Figures 4E-H). Reintroduction of full-length MAVS into the KO background restored antiviral control, reducing viral titers across multiple cellular systems (Figures 4I-K, Figures S4C&D). Notably, loss of MAVS also enhanced viral replication in primary MEFs and A549 cells – two systems with relatively low ACE2 expression (Figure S4E) and limited baseline permissivity to SARS-CoV-2(*48, 49*) – to levels comparable to those observed in highly permissive models (Figures 4L&M, Figures S4F&G). Collectively, these results suggest that MAVS is a major determinant of SARS-CoV-2 tropism and possibly a broader viral permissivity.

**Figure 4:**
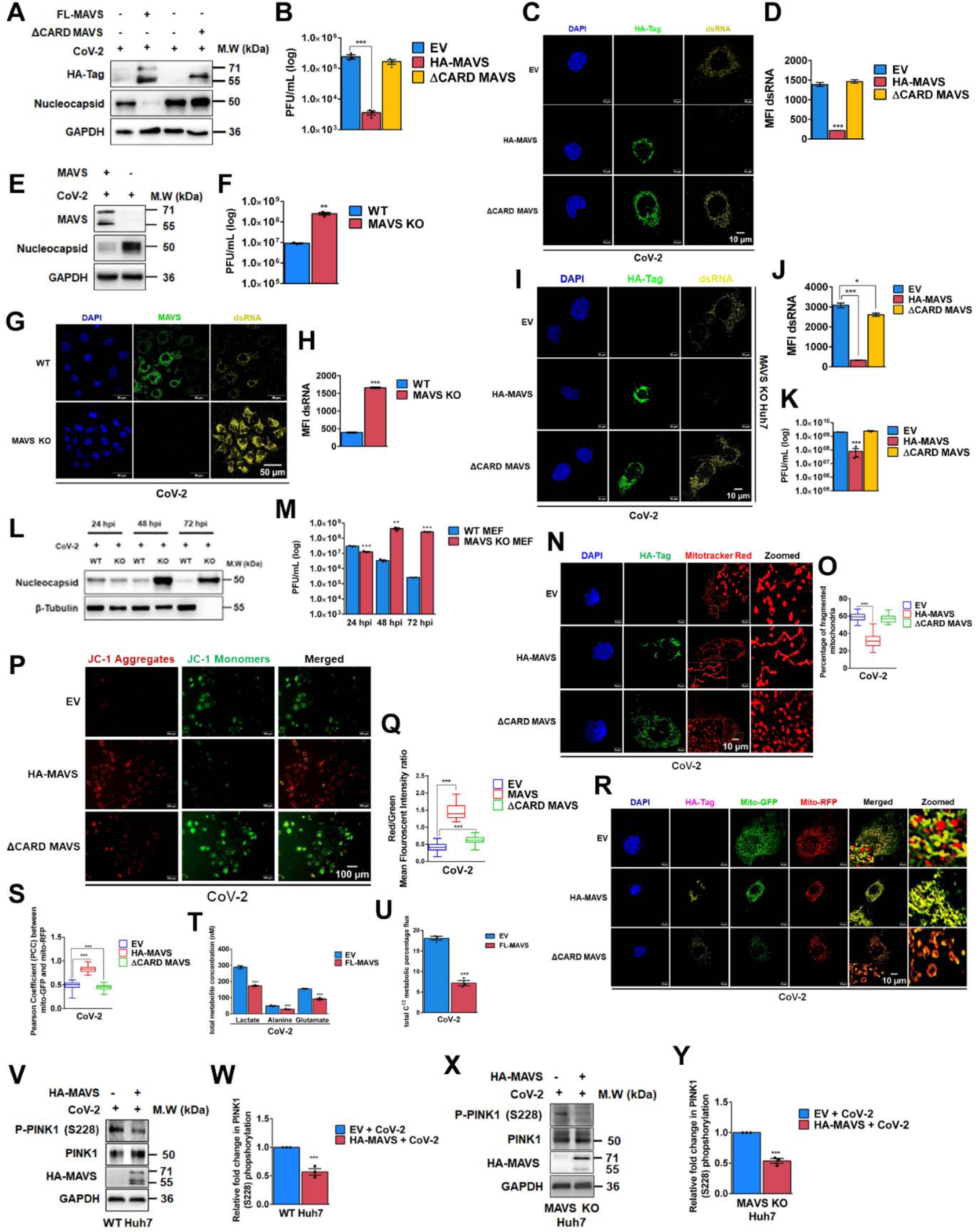
MAVS implements mitochondrial homeostasis to restrict RNA virus replication. **(A)** Immunoblot showing HA-MAVS and SARS-CoV-2 nucleocapsid levels in Huh7 cells overexpressing HA-tagged FL-MAVS or ΔCARD MAVS after SARS-CoV-2 infection. **(B)** Infectious viral titers from corresponding supernatants. **(C)** Confocal images detecting dsRNA in similarly treated cells, and **(D)** quantification of dsRNA mean fluorescence intensity (MFI). **(E)** Immunoblot showing MAVS and SARS-CoV-2 nucleocapsid levels in WT and MAVS KO Huh7 cells following infection, and **(F)** corresponding infectious viral titers. **(G)** Confocal images of dsRNA staining and MAVS expression in MAVS KO Huh7 and their WT cells **(H)** quantification of dsRNA intensity in samples from (G) (n = 100 cells). **(I)** Confocal images showing dsRNA in MAVS KO Huh7 cells transfected with HA-tagged FL-MAVS or ΔCARD MAVS and infected with SARS-CoV-2, and **(J)** dsRNA intensity quantification (n = 100 cells). **(K)** Infectious titers from corresponding supernatants. **(L)** Immunoblot analysing SARS-CoV-2 nucleocapsid levels in WT and MAVS KO MEFs infected with SARS-CoV-2. **(M)** Infectious viral titers comparing infection in WT and MAVS KO MEFs. **(N)** Confocal images analyzing mitochondrial fragmentation in SARS-CoV-2 infected Huh7 cells overexpressing HA-tagged FL-MAVS or ΔCARD-MAVS, and **(O)** corresponding quantification. **(P)** Epifluorescence microscopy images analyzing mitochondrial membrane potential in SARS-CoV-2 infected Huh7 cells upon supplementation of FL-MAVS or ΔCARD-MAVS. Scale bars = 100 µm, and **(Q)** Quantitative representation of mitochondrial membrane potential from images in (P) (n = 100 cells). **(R)** Confocal images analyzing mitophagy using mito-GFP-RFP assay in similarly treated cells, and **(S)** corresponding quantification. **(T)** Quantification of total glycolytic metabolite levels and **(U)** ATP synthesis rates by 1D metabolic NMR in SARS-CoV-2 infected Huh7 cells overexpressing FL-MAVS. **(V)** Immunoblot analysis of PINK1 phosphorylation in SARS-CoV-2 infected Huh7 WT cells overexpressing FL-MAVS, and **(W)** densitometric quantification. **(X)** Immunoblot analysis of PINK1 phosphorylation in SARS-CoV-2 infected MAVS KO Huh7 cells overexpressing FL-MAVS, and **(Y)** corresponding quantification. Scale bars, 10 µm and n = 50 cells unless otherwise indicated. Data are presented as mean ± SEM, with *, ** and *** denoting p <0.05, 0.01 and < 0.001.

Given MAVS’s role in mitochondrial homeostasis, we asked whether MAVS-mediated restriction preserves mitochondrial network organization during infection. Full-length MAVS, but not ΔCARD-MAVS, preserved mitochondrial network organization during SARS-CoV-2 infection, reducing infection-associated fragmentation (Figures 4N&O). Full-length MAVS, but not ΔCARD-MAVS, restored mitochondrial membrane potential and reduced infection-associated mitophagosome formation in SARS-CoV-2–infected cells (Figures 4P-S). MAVS supplementation produced similar rescue effects during JEV infection but not DENV infection, consistent with the limited mitophagy engagement in DENV infection (Figures S4H-K). Consistent with its role in maintaining mitochondrial integrity, MAVS supplementation in SARS-CoV-2–infected cells reduced glycolytic metabolite abundance and lowered ATP production rates, indicating a partial reversal of the virus-induced metabolic reprogramming (Figures 4T&U). MAVS supplementation also reduced infection-induced PINK1 accumulation in both WT and MAVS-KO cells across multiple systems (Figures 4V-Y, Figures S4L-N). Together, these data establish MAVS as a central mitochondrial safeguard that restricts viral replication by maintaining mitochondrial integrity and suppressing aberrant PINK1-dependent mitophagy during infection.

### MAVS restricts viral infection through an interferon-independent intrinsic antiviral program

We noticed with surprise that despite causing a potent SARS-CoV-2 suppression, MAVS supplementation during SARS-CoV-2 infection did not enhance IRF3 phosphorylation (Figures 5A&B; Figures S5A&B) nor induce *IFNB1* transcripts or secreted type I IFN (Figures 5C&D; Figure S5C). Importantly, MAVS robustly activated IRF3 phosphorylation and *IFNB1* expression and type I IFN in uninfected cells (Figures S5D-G), confirming intact interferon signaling competence in these cells. As a comparator, MAVS supplementation reduced JEV and DENV replication while enhancing type I IFN signaling (Figure S5H-M), consistent with canonical RLR-IRF3-IFN antiviral signaling in these IFN-inducing infections(*46, 50, 51*). By contrast, MAVS restricted SARS-CoV-2 replication without detectable IFN amplification, indicating a distinct interferon-independent MAVS activity in this context. Because peroxisomal MAVS can also drive type III interferon responses(*7*), we examined IFN-λ induction following MAVS supplementation. MAVS robustly induced *IFNL1* expression in both SARS-CoV-2 infected and uninfected cells (Figure 5E, Figure S5N). However, exogenous IFN-λ treatment failed to suppress SARS-CoV-2 replication (Figures 5F-H), indicating that type III interferon signaling does not account for MAVS-mediated restriction. RNA-seq showed that MAVS supplementation elicits a transcriptional program distinct from IFN treatment, most prominently during SARS-CoV-2 infection (Figure S5O), consistent with an IFN-independent antiviral state. Overlap analysis showed that MAVS and IFN share a comparable core ISG module in both mock and SARS-CoV-2 infection (∼63% overlap), while each condition also induced a distinct set of additional ISGs (Figures S5P&Q). Consistently, heatmaps revealed distinct MAVS- versus IFN-driven ISG signatures with a limited shared core (Figure S5R &S), supporting the possibility of involvement of a MAVS-driven antiviral program outside the canonical IFN-mediated during SARS-CoV-2 infection.

**Figure 5:**
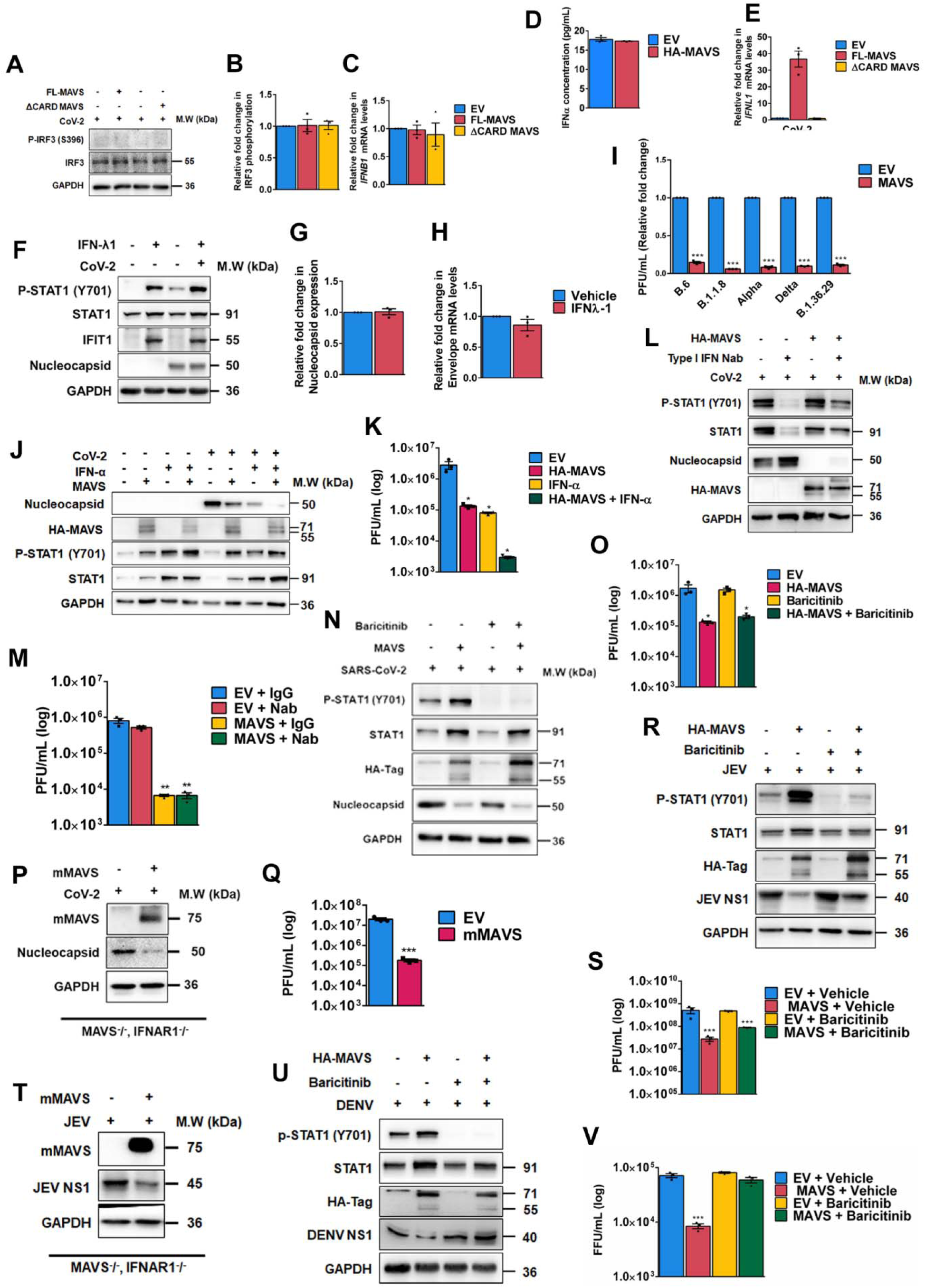
MAVS restricts viral infection through an interferon-independent intrinsic antiviral program. **(A)** Immunoblot analyzing IRF3 phosphorylation in SARS-CoV-2 infected Huh7 cells overexpressing FL-MAVS or ΔCARD MAVS. **(B)** Densitometric analysis of IRF3 phosphorylation. **(C)** qRT-PCR analysis of *IFNB1* mRNA levels and **(D)** ELISA for IFN-α from their corresponding supernatants. **(E)** qRT-PCR analysis of *IFNL1* mRNA levels from (A). **(F)** Immunoblot showing STAT1 phosphorylation, SARS-CoV-2 nucleocapsid, and HA-MAVS levels in IFN-λ-treated, SARS-CoV-2-infected Caco2 cells, and **(G)** nucleocapsid expression levels. **(H)** qRT-PCR of SARS-CoV-2 RNA from (F). **(I)** Relative fold change in infectious viral titers upon MAVS overexpression in Huh7 cells infected with different SARS-CoV-2 variants. **(J)** Immunoblot analyzing STAT1 phosphorylation, SARS-CoV-2 nucleocapsid, and HA-MAVS levels in SARS-CoV-2 infected Huh7 cells overexpressing MAVS, treated with IFN-α, or both, and **(K)** infectious viral titers from corresponding supernatants. **(L)** Immunoblot showing STAT1 phosphorylation, HA-MAVS, and nucleocapsid levels in Huh7 cells treated with type I IFN neutralizing antibodies, and **(M)** viral titers from their supernatants. **(N)** Immunoblot showing STAT1 phosphorylation, HA-MAVS, and nucleocapsid levels in Huh7 cells overexpressing MAVS, treated with Baricitinib, or their combination, and **(O)** corresponding viral titers from their supernatants. **(P)** Immunoblot showing mMAVS and nucleocapsid expression in SARS-CoV-2 infected MAVS:IFNAR DKO MEFs, and **(Q)** infectious viral titers. **(R)** Immunoblot analysis of STAT1 phosphorylation, HA-MAVS, and JEV NS1 expression in Huh7 cells overexpressing HA-MAVS, or treated with Baricitinib or their combination, upon infection with JEV, and **(S)** infectious viral titers from their supernatants. **(T)** Immunoblot analysis showing mMAVS and JEV NS1 expression in MAVS:IFNAR DKO MEFs overexpressing mMAVS upon JEV infection. **(U)** Immunoblot analysis of STAT1 phosphorylation, HA-MAVS, and DENV NS1 expression in Huh7 cells overexpressing HA-MAVS, or treated with Baricitinib or their combination, upon infection with DENV; and **(V)** foci formation assay quantification of infectious viral titers from their supernatants. For studies involving transfection and infection, cells were transfected with 2 µg of each construct, infected with SARS-CoV-2 or JEV (MOI = 1) 24 hours post-transfection (hpt), and stained or harvested 24 hours post-infection. Data are presented as mean ± SEM, with *, ** and *** denoting p <0.05, 0.01 and < 0.001.

We have used an earlier variant isolate of SARS-CoV-2 throughout in this study. Previous studies from our lab had shown that the earlier variants of SARS-CoV-2 show sensitivity to IFN-mediated antiviral defense, whereas later emerged variants including Alpha and Delta showed remarkable resistance to it(*10*). Consistent with its proposed IFN-independent activity, MAVS supplementation also restricted replication of several SARS-CoV-2 variants, including Alpha and Delta (Figure 5I). This indicated that MAVS-mediated restriction extends across SARS-CoV-2 variants that retain enhanced IFN evasion capabilities. MAVS supplementation and IFN-α treatment individually suppressed SARS-CoV-2 replication to comparable magnitudes. In support of MAVS’ IFN-independent action, their combination generated substantially profound reduction in viral titers (Figures 5J&K) indicating that they operated through parallel antiviral pathways. Further supporting these observations, MAVS-mediated suppression of SARS-CoV-2 replication prevailed unperturbed even under type I IFN receptor neutralization (Figures 5L&M), and pharmacological inhibition of JAK-STAT signaling by baricitinib (Figures 5 N&O). We generated MAVS/IFNAR1 double-knockout mice and tested MAVS’s antiviral potential in primary MEFs generated from them. MAVS supplementation significantly suppressed SARS-CoV-2 replication (Figures 5P&Q), confirming the interferon-independent MAVS antiviral mechanism in a genetic background. Because MAVS can also activate NF-κB(*6*), we inhibited NF-κB signaling with BAY11-7082 and found unaltered antiviral activity upon MAVS supplementation (Figure S5T&U), indicating that MAVS-mediated SARS-CoV-2 restriction is also NF-κB–independent. Similarly, when interferon signaling was blocked pharmacologically or genetically, MAVS supplementation continued to suppress JEV replication. This result suggested that the IFN-independent viral restriction operates in parallel to the IFN-mediated mechanisms, and that it can continue to operate even upon IFN-suppression (Figures 5R-T). In contrast, MAVS-mediated suppression of DENV reverted upon interferon blockade, indicating that the IFN-independent antiviral defense operating upon SARS-CoV-2 and JEV infections do not operate during DENV infection. This is consistent with the clear absence of mitophagy during DENV infection and the lack of impact of mitophagy on its replication (Figures 5U&V). Collectively, these data establish that MAVS elicits antiviral response that is not restricted through IFN- and NF-κB signaling outputs, but also by invoking a very potent intrinsic alternate mechanism that involves mitochondrial stabilization.

### MAVS-mediated mitochondrial safeguarding is not contingent upon IFN signaling

Finally, to verify if the MAVS-dependent intrinsic antiviral activity involves mitochondrial remodeling independent of canonical signaling outputs, we monitored mitochondrial responses to RNA and interferon stimulation. We studied this in a set up using Contact-ID, a split-BioID reporter system based on a TOM20–Sec61 pair that labels mitochondria in the context of mitochondria–ER junctions (*52*). Poly(I:C) pronounced mitochondrial elongation in WT cells, whereas MAVS-KO cells failed to remodel mitochondrial networks in response to RNA sensing (Figures 6A&B). Similarly, IFN-α treatment elongated mitochondria in WT Huh7 cells but had no effect in the absence of MAVS (Figures 6C&D). Given that mitochondrial elongation has been linked to enhanced mitochondria–ER communication(*53, 54*), we next assessed mitochondria–ER contact sites (MAMs). In line with our mitochondrial elongation results, Poly(I:C) treatment in WT cells induced a strong increase in MAM/MERCs, but failed to do so in MAVS KO cells (Figures S6A-C). Together, these data indicate that MAVS is critical for both RNA- and IFN-triggered mitochondrial remodelling.

**Figure 6:**
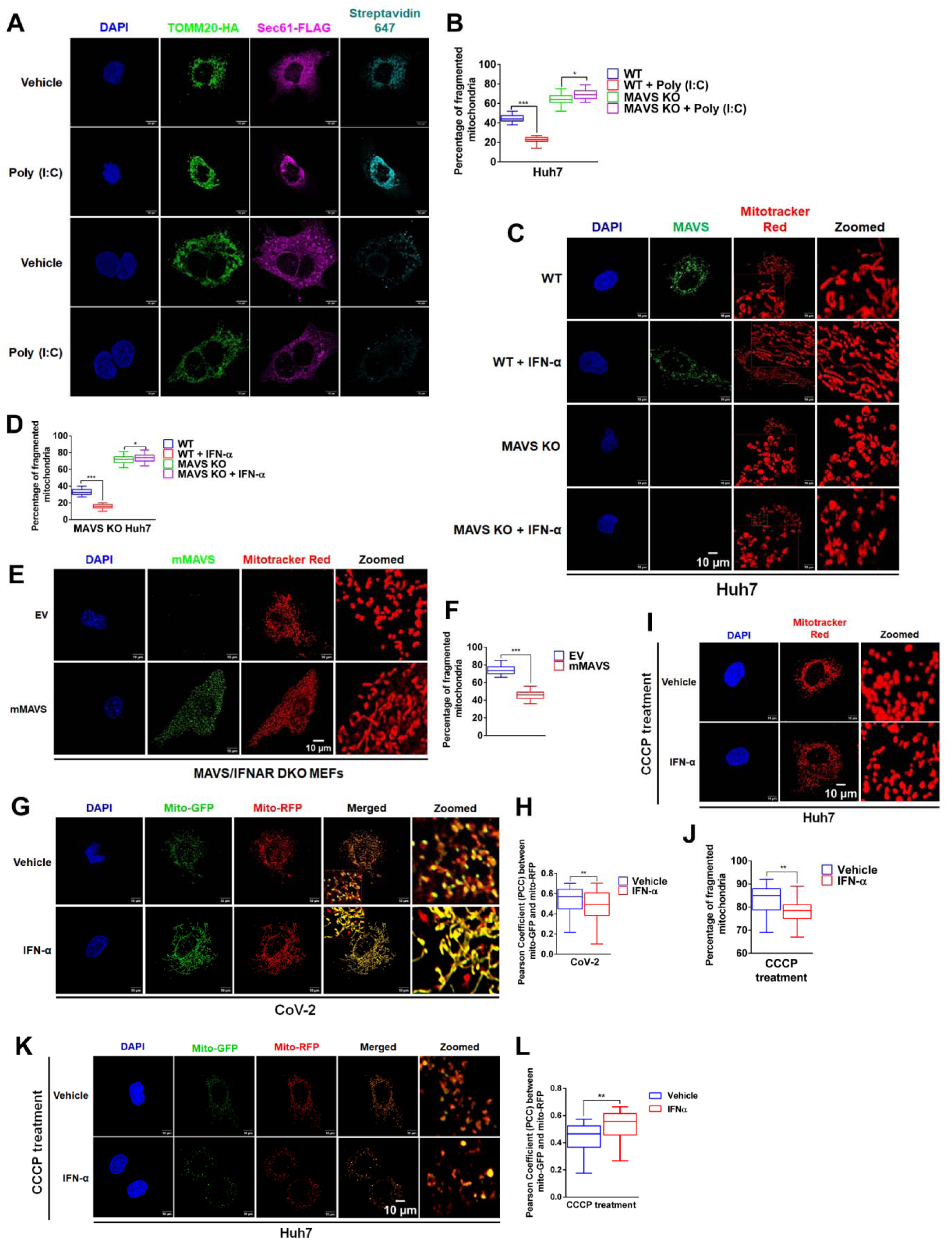
MAVS-mediated mitochondrial safeguarding is not contingent upon IFN signaling. **(A)** Super-resolution images showing HA-tagged TOM20 (mitochondria) and FLAG-tagged Sec61 (ER), along with their contact sites (MAM) in Vehicle or Poly(I:C) treated WT and MAVS KO cells. **(B)** Quantification of mitochondrial fragmentation from experimental setup described in (A). **(C)** Confocal images analyzing mitochondrial fragmentation in IFN-α treated Huh7 WT and MAVS KO cells, and **(D)** corresponding quantification. **(E)** Confocal images analyzing mitochondrial fragmentation in MAVS/IFNAR DKO MEFs overexpressing mMAVS, and **(F)** corresponding quantification. **(G)** Confocal images analyzing mitophagy in CoV-2 infected, IFN-α treated Huh7 cells, and **(H)** corresponding quantification. **(I)** Confocal images studying mitochondrial fragmentation in CCCP and IFN-α treated Huh7 cells, and **(J)** Quantification of mitochondrial fragmentation from (I). **(K)** Confocal images showing mitophagy in CCCP and IFN-α treated Huh7 cells, and **(L)** Quantification of mitophagy from images in (K). Scale bars, 10 µm and n = 50 cells unless otherwise indicated. Data are presented as mean ± SEM, with *, ** and *** denoting p <0.05, 0.01 and < 0.001.

We further validated this IFN-independence in MAVS/IFNAR1 double-knockout cells. MAVS supplementation restored mitochondrial network organization in this IFN-deficient background (Figures 6E&F). Further, IFN-α treatment during SARS-CoV-2 infection failed to restore either mitochondrial membrane potential (Figures S6D&E), or reduce mitophagosome accumulation (Figures 6G&H) to the extent as observed with MAVS supplementation (Figures 4P-S). To determine whether interferon can broadly counteract mitochondrial damage independent of viral infection, we examined mitochondrial morphology following CCCP-induced depolarization. IFN-α treatment failed to prevent CCCP-induced mitochondrial fragmentation (Figures 6I&J), membrane depolarization (Figures S6F&G), and mitophagosome accumulation (Figures 6K&L), indicating that interferon signaling is insufficient to suppress mitochondrial damage even in a non-viral context. These results confirm that the mitochondrial stabilization by MAVS is completely independent of IFN-signaling activities. Collectively, these findings establish MAVS as a regulator of intrinsic antiviral immunity that preserves mitochondrial integrity and import competence, thereby restricting viral replication independently of canonical interferon and NF-κB signaling pathways.

## DISCUSSION

Antiviral defense is layered: in addition to inducible innate immune signaling that culminates in interferons and cytokines, cells deploy intrinsic, cell-autonomous defenses that restrict infection by maintaining a non-permissive intracellular state. Intrinsic immunity is characterized by specialized host restriction factors that employ discrete mechanisms to target specific stages of the viral life cycle. But are there any unifying themes to the intrinsic antiviral immunity? Our findings suggest so. Here, we show that mitochondria play crucial roles in enacting a potent intrinsic immunity independent of IFN. MAVS, previously known for its role in coordinating IFN production, is a decisive player in the intrinsic immunity as well. We show that MAVS orchestrates this by safeguarding mitochondrial integrity through ensuring proper mitochondrial protein import and thereby restraining PINK1-driven mitophagy. This IFN-independent mitochondrial safeguard, exploited by RNA viruses (like SARS-CoV-2 or JEV), enforces viral restriction even when canonical IFN outputs are weak or blocked. Given that the RLR pathway is also critically dependent on mitochondrial integrity for signalosome complex assembly, our findings suggest that MAVS’s primary function is the safeguarding of mitochondria and its antiviral functions are secondary outcomes of this function (Figure 7).

**Figure 7:**
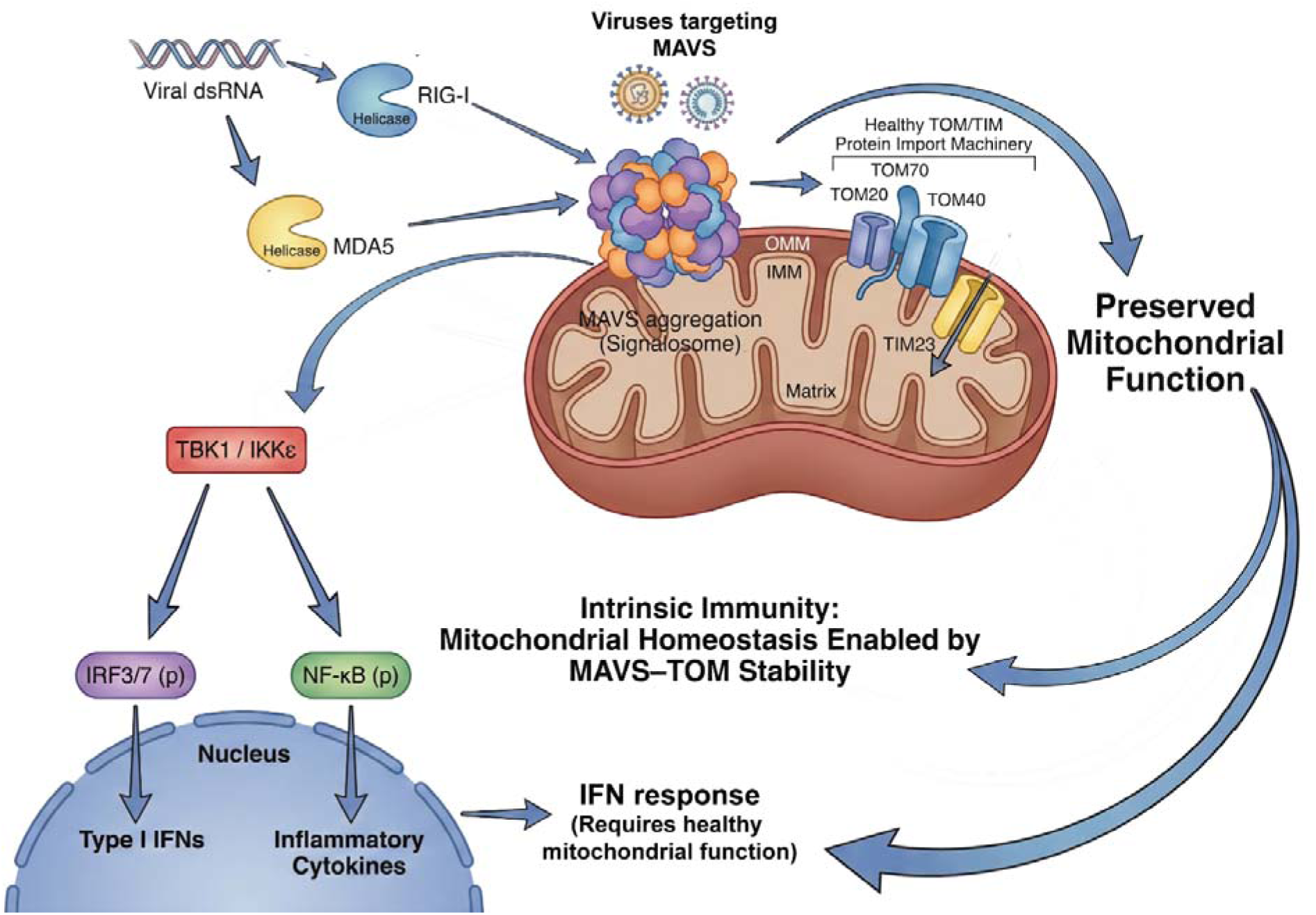
MAVS-dependent antiviral signaling is coupled to mitochondrial protein import and organelle homeostasis. Viral RNA sensing by RIG-I and MDA5 activates MAVS on mitochondria, triggering TBK1/IKKε signaling and induction of type I interferons and inflammatory cytokines. MAVS function is coupled to the TOM/TIM protein import machinery, preserving mitochondrial integrity required for effective antiviral responses.

MAVS possesses an extensive interactome map that shows its potential interactions with over 250 proteins(*55*). Of these, interactions with TOM20 and MFN1/2 indicated a potential influence by this molecule on mitochondrial homeostasis. Indeed, our deep-sequencing indicated strongly in that direction. We demonstrate that MAVS is critical for proper assembly of TOM/TIM complex. Mechanistically, our data supports a model in which MAVS preserves the integrity of the mitochondrial import apparatus and its higher-order organization on the outer membrane. Loss of MAVS selectively depletes key TOM/TIM receptor and channel components, perturbs TOM/TIM assemblies, and compromises import, which are defects that are expected to destabilize respiratory function and membrane potential. Import failure provides a direct route to PINK1 stabilization and engagement of mitophagy, linking MAVS-dependent mitochondrial competence to suppression of PINK1-dependent quality control programs that otherwise remodel (and potentially deplete) the mitochondrial network.

Many RNA viruses antagonize MAVS to suppress antiviral signaling; our findings indicate that this interference carries a second, underappreciated consequence, which is destabilization of mitochondrial homeostasis itself. During SARS-CoV-2 infection, MAVS fails to undergo productive aggregation, coinciding with mitochondrial fragmentation, loss of membrane potential, PINK1 stabilization, and elevated mitophagosome formation. Our results indicate that these changes are not unique to SARS-CoV-2 infection, but represent many such viruses that promote mitophagy, as we shown in JEV. These defects closely phenocopy MAVS deficiency, suggesting that viral disruption of MAVS assembly simultaneously dampens canonical antiviral signaling and drives mitochondria into an import-compromised, quality-control–engaged state. Together, this provides a tight mechanistic link between viral immune evasion at the level of MAVS and infection-associated mitochondrial dysfunction.

A key implication of our findings is that mitophagy should not be viewed solely as a source of membrane platforms for viral replication. Instead, loss of mitochondrial mass and integrity can itself erode antiviral competence, particularly in non-immune cells where mitochondrial fitness underpins both signaling capacity and metabolic flexibility. Consistent with this view, experimentally inducing mitophagy enhanced replication of viruses that benefit from mitochondrial pruning, such as SARS-CoV-2 and JEV, whereas viruses that rely on intact mitochondrial function or employ distinct remodeling strategies, exemplified by DENV, did not benefit and were instead impaired. These observations underscore that the consequences of mitophagy are virus-specific and tightly coupled to how individual pathogens exploit or tolerate mitochondrial dysfunction.

Conceptually, our study illustrates that mitochondrial integrity serves as a master regulator of multi-layered innate immune responses. The mitochondrial regulatory landscape—encompassing both structural maintenance and metabolic homeostasis—is intrinsically coupled to the cell’s capacity for broad-spectrum antiviral defense. Specifically, we highlight how fundamental processes, such as mitochondrial protein import, function as critical determinants of the cellular antiviral state. Notably, the modulation of the expression of TOM complex components through transcriptional regulation by MAVS suggested that MAVS exerts far-reaching control over mitochondrial architecture. Given the extensive inter-organellar communication networks, these MAVS-dependent mitochondrial alterations likely reverberate throughout the cell, impacting the functional homeostasis of associated organelles. The evolutionary pressure on viruses to develop sophisticated strategies for subverting MAVS further underscores its pivotal role in maintaining the host environment. Ultimately, by unraveling these multifaceted regulatory nodes, our study significantly advances the conceptual framework of the innate antiviral response.

“Intrinsic immunity” has traditionally been defined as constitutive, cell-autonomous restriction mediated by pre-existing antiviral factors. Our findings extend this concept by identifying organelle fitness as a major determinant of intrinsic antiviral competence, with MAVS acting as a regulator of mitochondrial integrity at the organelle surface. This role may be particularly consequential in non-immune cell types such as epithelial cells and fibroblasts, where preservation of mitochondrial networks and mitochondria–ER contact sites can determine the baseline capacity for antiviral signaling and metabolic adaptation, even before cytokine amplification occurs. An additional implication of our data is that mitochondrial safeguarding by MAVS may indirectly modulate inflammatory outcomes. Severe mitochondrial perturbation can promote danger signaling, including mtDNA release and inflammasome activation. However, the primary phenotypes we observe – import failure, loss of membrane potential, PINK1 stabilization, and engagement of mitophagy – arise upstream of overt inflammatory cell death. We therefore favor a model in which MAVS-dependent mitochondrial integrity restrains excessive quality-control activation and secondary danger signaling by preventing escalation of mitochondrial damage, a function that may be especially relevant in epithelial immune circuits during viral infection.

Two limitations of this study merit consideration. First, MAVS supplementation likely “preconditions” cells, and the extent to which MAVS can be therapeutically restored after infection onset may be constrained by virus-induced stress or cytotoxicity. Second, the precise molecular mechanism by which MAVS stabilizes mitochondrial import machinery remains unresolved. MAVS may act through direct scaffolding of TOM/TIM assemblies, by organizing outer-membrane and MERC architecture, or indirectly by preserving membrane potential and mitochondrial ultrastructure. Future work should delineate which MAVS domains beyond the CARD region are required for TOM receptor clustering and import competence, identify viral factors responsible for MAVS puncta depletion or redistribution, and test whether genetic or pharmacologic inhibition of PINK1-dependent quality control can restore antiviral competence downstream of MAVS loss.

Together, our findings elevate MAVS from a mere signal-transduction adaptor to a higher order position of a key mitochondrial guardian. Its unique position also exposes MAVS as a handle for the viruses on the host antiviral immunity where targeting it provides an overarching control over the antiviral response. This organelle-centric function of MAVS provides a mechanistic bridge between viral immune evasion and mitochondrial remodeling, and suggests that strategies aimed at preserving mitochondrial fitness may offer therapeutic leverage even when canonical interferon pathways are compromised.

## MATERIALS AND METHODS

### Antibodies

MDA-5 (Cat # 5321), RIG-I (Cat #3743), Human MAVS (Cat #24930), Rodent MAVS (Cat #4983), Phospho IRF-3 (Cat #4947), IRF3 (Cat #11904), Phospho-STAT1 (Cat #14994), STAT1 (Cat #9167), IFIT1 (Cat #14769), Phospho P65 (Cat #3033), P65 (Cat #8242), HA-tag (Cat #3724 and #2367), FLAG-tag (Cat#14793), LC3 A/B (Cat #12741), TOM70 (Cat #65619), TOM20 (Cat #42406), TOM22 (Cat# 90704), TOM40 (Cat# 55959) were purchased from Cell Signaling Technologies (CST). TIM23 (Cat #sc-514463) and TOM40 (Cat #sc-365467) was from Santa Cruz Biotechnology. JEV NS1 (Cat #GT1410) and DENV NS1 (Cat #GTX124280) antibodies were from Genetex. dsRNA J2 antibody (Cat #10010200) was procured from Scicons. Phospho PINK1 (Cat #PA5-105356), PINK1 (Cat #PA5-86941), CoV-2 nucleocapsid (Cat #MA5-36272), GAPDH (Cat #MA5-15738), β-Tubulin (Cat #MA5-16308), ACE2 (Cat #MA5-32307), Alexa Flour 647 conjugated with Streptavidin (CAT#S21374 were procured from Thermo Fisher Scientific. Goat Anti-rabbit HRP conjugated (Cat #115-035-146), Alexa Flour 488 conjugated (Cat #115-545-146), Alexa Four 647 conjugated (Cat #115-605-146) and Goat anti-mouse HRP conjugated (Cat #115-035-003), Alexa Flour 488 conjugated (Cat #115-545-003), Alexa Flour 594 conjugated (Cat #111-585-006) and Alexa Flour 647 conjugated (Cat #115-605-003) were purchased from Jakson ImmunoResearch.

### Inhibitors and reagents

Human IFNα 2a (Cat #11100) was purchased from PBL assay science. Poly (I:C) (Cat #P1530), BAY-11-7082 (NF-κB inhibitor, Cat #196870), Baricitinib (JAK-STAT inhibitor, Cat #SBR00076) and Carbonyl Cyanide m-Chlorophenylhydrazone (CCCP, Cat #C2759) was from Sigma-Aldrich. Human Type I IFN Neutralizing Antibody Mixture (Cat #39000) was from PBL assay science. Human IFN-lambda was from R & D systems (Cat # 1598-IL) Human IFN alpha ELISA Kit (Cat #BMS216) and JC-1 dye (Cat #T3168) was from Invitrogen. Mitotracker Red (CMX-ROS) dye (Cat #9082) was from Cell Signaling Technologies (CST).

### Plasmids and cloning

Mammalian expression vector, pcDNA4/TO was from Invitrogen. Mammalian expression vector pUNO and HA-MAVS were from Invivogen. HA-ΔCARD-MAVS were cloned as reported previously(*38*). Mouse MAVS was amplified using cDNA of MEFs and cloned into pcDNA4/TO. Mitophagic flux was assessed by using a p-mito-mRFP-EGFP plasmid (Plasmid pAT016)(*56*). Mitochondrial protein import assay was performed using a MTS-GFP fusion protein cloned in pcDNA/4TO plasmid(*57*). MAM formation was assessed using Contact-ID – a split-BioID system(*52*).

### Generation of MAVS and IFNAR1 Double KO MEFs

All mice were bred and maintained under specific pathogen-free conditions at the CCMB Animal Facility. Double knockout mice for MAVS and IFNAR1 were generated using MAVS KO males (B6129SF2/J, #008634, Jackson Laboratory) and IFNAR1 KO females (C57BL/6J, #028288). Appropriate animal ethical clearances were obtained from the Institutional Animal Ethics Committee of CCMB and their guidelines were strictly followed. First-generation (G1) heterozygous offspring (MAVS +/-; IFNAR1 +/-) were genotyped and interbred to produce G2 mice, including double homozygotes (MAVS -/-; IFNAR1 -/-). Genomic DNA was prepared from their tail samples (1–2 mm). Were genotyped by PCR using gene-specific primers targeting wild-type and mutant alleles. Amplicons were analyzed via agarose gel electrophoresis. MAVS KO yielded a 350 bp band, IFNAR1 KO a ∼450 bp band, and heterozygotes showed both bands. Thermal cycling conditions were optimized separately for each gene.

### Cell culture

Caco2 cells were obtained from ATCC, while Huh7 cells were generously provided by Ralf Bartenschlager(*58*). A549 WT and MAVS KO cells were from Invivogen. MEFs were isolated from WT, MAVS KO and MAVS:IFNAR1 DKO mice following the established protocol(*59*). All cells, except for Caco2 (20% FBS), were maintained in DMEM (Gibco) supplemented with 10% FBS (Gibco), 1× penicillin-streptomycin cocktail (Gibco), and 1× non-essential amino acids (Gibco) at 37°C with 5% CO2. Cells were routinely passaged at 70–80% confluency, and mycoplasma contamination was periodically monitored.

### Generation of MAVS KO Huh7

MAVS KO Huh7 cells were generated using the CRISPR-Cas9 system. Single-guide RNAs (sgRNAs) targeting exon 6 and 8 of the MAVS gene inserted into the PX459 vector containing WT Cas9 (Addgene, Cat #62988). Huh7 cells were transfected with the sgRNA-containing PX459 plasmids using Lipofectamine 3000. Following transfection, puromycin selection was used to enrich transfected cells. Knockout validation was performed by immunoblotting in multiple clones to confirm the loss of MAVS protein. Successfully edited clones were expanded and verified as MAVS knockout Huh7 cells. The following sgRNA primers were used:

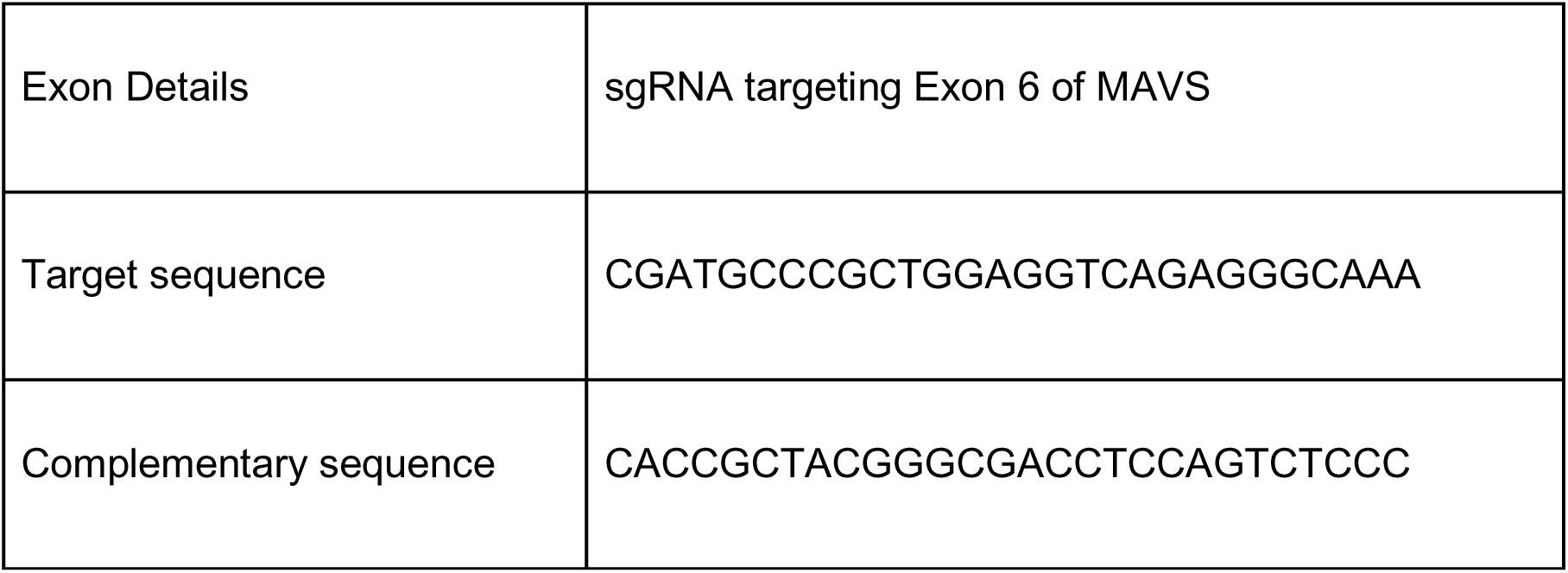

### JEV, DENV and SARS-CoV-2 isolates

The five variant isolates of SARS-CoV-2 used in this study were isolated at the Centre for Cellular and Molecular Biology in the biosafety level 3 facility(*10*). The viruses were propagated in Vero (CCL-81) cells grown in serum free DMEM. JEV and DENV were propagated as previously described(*38*).

### Virus Infection and quantification

Cells were infected at 1 MOI for 3 hrs in serum-free conditions after which the media was replaced with complete media and further incubated until the time of harvesting as per experiment requirement. During harvesting, the cells were first trypsinized and collected separately for protein and RNA study. Titration for SARS-CoV-2, JEV and DENV were done as described(*10, 38*). For Overexpression studies, cells were seeded to reach 50 to 60% confluency. Transfection mix containing Opti-MEM, Lipofectamine 3000, along with plasmid of interest was prepared according to the manufacturer’s protocol, added to cells, and incubated for 6 h. Later, the transfection mix was replaced with cDMEM and further incubated for 18 h. 24 hours post transfection, cells were infected with virus for 3 h and further incubated in fresh cDMEM until they were harvested for analyses. Similar approach was used for Poly(I:C) transfections.

For IFN-α and IFNλ-1 treatment, cells were serum-starved at 80 to 85% confluency in serum-free DMEM (SFD) for 3 h. Later, SFD was replaced with fresh SFD containing 1000 U/mL IFN-α or IFNλ-1 for 1 h. Following this, the cells were infected for 3 h, and incubated further with fresh cDMEM containing either phosphate-buffered saline (PBS; vehicle) or 1,000 U/mL of IFN-α or IFNλ-1, and incubated for the subsequent 21 h. Cells and culture supernatants were harvested at 24 hpi and used for further analysis.

For Baricitinib and BAY-11-7082 treatments, Cells were treated with the inhibitors 3 hpi and incubated for the subsequent 21 h. For CCCP treatments, cell culture media was removed and cells were supplemented with CCCP containing cDMEM 2 hours and 6 hours before harvesting, respectively. Cells and Cell culture supernatants were harvested and used for further analysis.

### Real-Time quantitative RT-PCR (qRT-PCR)

Cellular RNA samples were prepared using an MN NucleoSpin RNA kit (TaKaRa, Cat #740902). Equal quantities of RNA were reverse transcribed using Primescript 1st strand cDNA synthesis kit (TaKaRa, Cat #6110) following the manufacturer’s protocol. 50□ng of cDNA was used for quantification using TB Green Premix (TaKaRa, Cat #RR820) on a LightCycler 480 instrument (Roche). Transcripts of the host origin were normalized against GAPDH. qRT-PCR to quantify SARS-CoV-2 RNA was performed on a Roche LightCycler 480 using an nCOV-19 reverse transcription-PCR (RT-PCR) detection kit from Q-Line Molecular (Cat #COVIDM96PS). Relative fold changes between the experimental and control samples (2–ΔΔCT) were calculated and represented in the graphs. Primer sequences for *IFNB1* and *DENV* are reported earlier(*38*). The following primers were used for qRT-PCR of *IFNL1*:

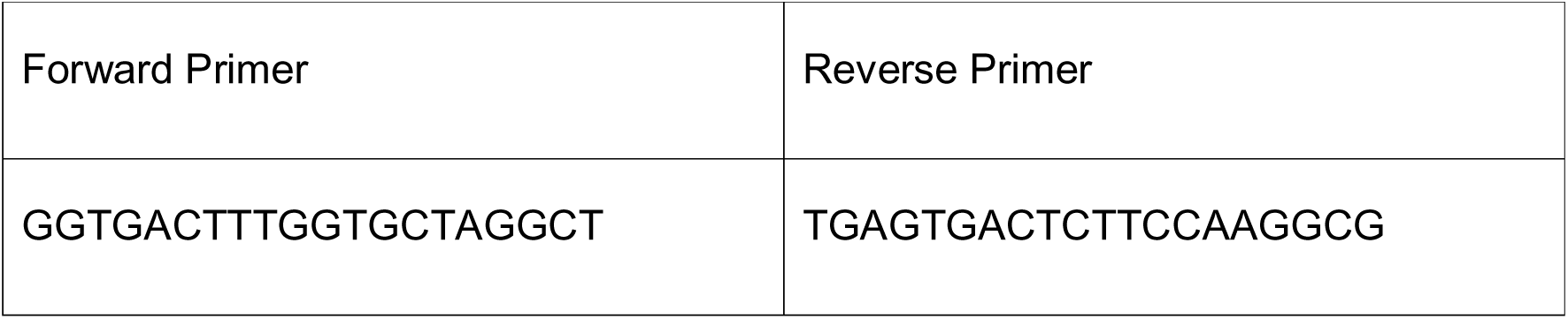

### Immunoblotting

Cell pellets were lysed in 1 × Nonidet P-40 lysis buffer (1% Nonidet P-40, 50 mM Tris-HCl, 150 mM NaCl (pH 7.5), EGTA, 1 mM sodium orthovanadate, 10 mM sodium pyrophosphate, 100 mM NaF, and 1 mM PMSF), incubated on ice for 20 minutes with intermittent vortexing and centrifuged at 13,000 rpm for 15 minutes at 4°C. The supernatants containing the proteins were collected and quantified using BCA reagents (Pierce, Cat #23225). Lysates were mixed with 6× denaturing dye and the proteins were resolved using SDS-PAGE and transferred to PVDF membranes. The membranes were blocked in 5% BSA dissolved in 1× TBST before the addition of primary antibodies. Primary antibodies against the proteins of interest were diluted in the blocking buffer, added to the membrane, and incubated overnight at 4°C. Later, the membranes were washed in 1× TBST, secondary antibodies conjugated with HRP were added and the blots were developed on a Bio-Rad Chemidoc MP system using Clarity Western ECL substrate (Biorad, Cat #1705061) and Clarity Max Western ECL substrate (Biorad, Cat #1705062) chemiluminescent substrate kits. Relative fold changes in protein expression between the experimental and control samples were determined and depicted in the graphs.

### Blue Native-PAGE

To assess TOM complex stability and abundance, Blue Native PAGE (BN-PAGE) was performed following established protocols(*60*). Briefly, Protein content in mitochondria-enriched fractions was quantified using Bradford Assay reagent, then samples were solubilized as described previously: digitonin (Sigma-Aldrich – Cat# D141-500MG) was added at 12 g per g of mitochondria-enriched fraction and incubated 10 min on ice, followed by 30 min centrifugation at 13,000 g to collect the supernatant. The protein concentration of this supernatant was re-measured by Bradford, then 5% G-250 sample dye was added at 6 g per g of mitochondrial-soluble fraction, and glycerol was adjusted to a final 10%.

20–30 µg of sample was loaded onto 4–15% Mini-PROTEAN® TGX™ Precast Protein Gels (Biorad – Cat#4561086) and run with anode buffer (50 mM Bis-Tris HCl, pH 7.0) and blue cathode buffer (50 mM Tricine, 15 mM Bis-Tris HCl, pH 7.0, 0.02% Coomassie Brilliant Blue G-250). Electrophoresis was performed at 100 V until entry into the separating gel, then 150 V until the dye front reached ∼1/3 of the gel, after which the blue cathode buffer was replaced with dye-free cathode buffer and the run continued at 250 V until the dye front reached the gel end.

### Seahorse Mito stress assay

Mitochondrial oxygen consumption was assessed using a Seahorse XFe 24 Analyzer. In summary, 50,000 cells per well were plated in XFe 24 plates, washed twice with DPBS, and incubated for 1 hour at 37 °C in a CO_₂_-free incubator with XFe DMEM assay medium containing 10 mM glucose, 2 mM L-glutamine, and 1 mM sodium pyruvate. Oligomycin (3.5 μM), FCCP (3 μM), and a combination of antimycin A (1.5 μM) with rotenone (1.5 μM) were introduced through reagent ports A, B, and C, respectively. Measurements were taken for basal oxygen consumption rate (OCR), ATP-linked OCR, and proton leak, with all values normalized to total protein content determined using the Bradford assay.

For overexpression studies, cells were transfected with the appropriate plasmids in 6-well plates for 24 hours, followed by trypsinization and reseeding at a density of 50,000 cells per well in XFe 24 plates. After reseeding, the same Seahorse assay procedure was carried out as previously described.

### Preparation and NMR Analysis of Metabolites from Cell Pellets

Cells incubated with medium containing [1,6-^13^C] glucose for 45 minutes were harvested and stored at -80°C for further analysis. Metabolites were then extracted from harvested cells using an ethanol-based extraction protocol(*61*). In brief, cells pellet was homogenized in methanol-HCl (0.1N) in (2:1 vol/wt) using a battery-operated tissue homogenizer. [2-^13^C] Glycine (50 μL, 2 mmol/L) was added as an internal reference, followed by homogenization with ethanol (90%)-phosphate buffer (6:1 vol/vol). The homogenate was centrifuged (20,000 g, 45 min, 4°C), and the supernatant pH-adjusted, and freeze-dried. The lyophilized powder was dissolved in phosphate buffer (25 mmol/L, pH 7.0, 80% D_2_O) containing sodium trimethylsilyl propionate (0.25 mmol/L) for NMR measurements. Total and ^13^C-labeled concentrations of metabolites were measured in ^1^H-[^13^C]-NMR spectrum at 600 MHz NMR spectrometer(*62*). The ^13^C-edited spectrum was obtained through spin-echo acquisition with and without a ^13^C inversion pulse. Metabolite levels were quantified relative to [2-^13^C] glycine, and ^13^C enrichment was corrected for natural abundance (subtracting 1.1%). The rate of glucose consumption (MR Glc) as calculated from the ^13^C labelled trapped into different metabolites(*63*). The rate of ATP synthesis was calculated as described previously using an approximation of 2.5 and 1.5 ATP yield for each molecule of NADH and FADH, respectively:

ATP syn = (2 NADH + 2 ATP) x Lac Syn + (10 NADH + 2 FADH 2 + 4 ATP) x MR Glc (Lac Syn is rate of lactate synthesis).

### Microscopy experiments

#### JC-1 Staining

50,000 cells were seeded in 6-well plates and allowed to adhere overnight. Cells were subjected to experimental conditions (virus infection, treatments, overexpression, or phenotype comparison), washed twice with SFD, and stained with 10 µM JC-1 dye in SFD for 20–30 minutes at 37°C in the dark. After staining, cells were washed twice with SFD, replenished with fresh SFD, and imaged using a Zeiss Axio Observer 3 microscope with FITC and mCherry filters to visualize green JC-1 monomers and red aggregates. ΔΨm was expressed as the JC-1 ratio = (Red_mean − Red_bg) / (Green_mean − Green_bg). Ratios were calculated per cell and summarized as mean ± SEM across biological replicates.

### Mitotracker Red staining

50,000 cells were seeded on coverslips in 6-well plates, subjected to experimental conditions, washed twice with SFD, and incubated with 100 nM Mitotracker Red or Mitotracker Green in SFD for 20–30 minutes at 37°C in the dark. After staining, cells were washed twice with SFD, fixed with 4% formaldehyde for 30 minutes, washed with PBS, and stored at 4°C before immunostaining.

### Mito-GFP-RFP Reporter Assay

50,000 cells were seeded on coverslips, transfected with 2 µg of Mito-GFP-RFP reporter plasmid, and subjected to experimental conditions 24 hours post transfection. Cells were washed twice with PBS, fixed with 4% formaldehyde for 30 minutes, washed with PBS, and stored at 4°C before immunostaining. Mitophagy was quantified as the Pearson correlation coefficient (PCC) between the GFP and RFP channels, with lower PCC values indicating increased mitophagy due to loss of GFP fluorescence in acidic lysosomal compartments while RFP signal is retained. PCC values were calculated per cell and averaged across cells and independent biological replicates.

### Mitochondrial protein import assay

50,000 cells were seeded on coverslips, transfected with 2 µg of MTS-GFP reporter plasmid and mCherry as a transfection normalization control. 12 hours post transfection, cells were washed twice with PBS, fixed with 4% formaldehyde for 30 minutes, washed with PBS, and immunostained. For WT + CCCP conditions, cells were transfected with MTS-GFP for 10 hours, followed by CCCP treatment for 2 hours before fixing the cells. For MAVS supplementation assays, empty vector, FL-MAVS and ΔCARD MAVS were co-transfected along with MTS-GFP and mCherry and were fixed 12 hours post-transfection before proceeding for immunostaining.

### MAM Formation Assay

50,000 cells were seeded on coverslips, co-transfected with 2 µg of HA-TOM20-BirA and FLAF-Sec671-BirA reporter plasmid pair (Contact ID). 24 hours post-transfection, cells were biotinylated with 50 μM of biotin and subjected to experimental conditions 24 hours post biotin treatment. Cells were washed twice with PBS, fixed with 4% formaldehyde for 30 minutes, washed with PBS, and stored at 4°C before immunostaining.

### Immunostaining

Fixed coverslips were washed twice with PBS, permeabilized (0.05% Tween-20, 0.25% Triton X-100 in PBS) for 10 minutes, and washed three times with PBST. Blocking was performed with 1% BSA and glycine in PBST. Coverslips were incubated overnight at 4°C with primary antibodies diluted in 1% BSA in PBST. After washing, fluorophore-conjugated secondary antibodies were added, followed by three washes with PBST. Coverslips were mounted with DAPI-containing anti-fade mountant (Cat #H-1200-10) and analyzed by microscopy.

### Imaging and Analysis

Confocal microscopy was performed using an Olympus FV3000 and Leica SP8 microscopes (60× oil immersion objective for Olympus FV3000, 63x oil immersion objective for Leica SP8, 2.5× optical zoom). Super-resolution imaging for contact-ID assay was conducted using a ZEISS Elyra 7 system with Lattice SIM² (63× objective, variable zoom). Mitochondrial fragmentation, volume analysis of clumped mitochondria (in SARS-CoV-2 and JEV-infected cells) were analyzed using Imaris software. JC-1 red/green ratios and co-localization (e.g., GFP-RFP, MAVS-mitochondria) were quantified with ImageJ. Binary HA-TOM20-FLAG-Sec61 intersections for MAMs were also calculated using ImageJ.

### Intersection density and spatial proximity analysis

Confocal images of Huh7 WT and MAVS KO cells were acquired using the Olympus FV3000 system. Post-acquisition, image stacks containing TOM20, TOM70, and Mitotracker Red channels were deconvolved using the DeconvolutionLab2 plugin in ImageJ, applying the Richardson–Lucy algorithm (20 iterations). Following deconvolution, TOM20 and TOM70 channels were merged, thresholded, and intersection masks were generated using ImageJ’s Image Calculator function. Intersections were quantified using the Analyse Particles tool. Mitochondrial area was determined from the Mitotracker Red channel, and intersection density was calculated as the number of TOM20–TOM70 intersections per unit mitochondrial area. For spatial proximity analysis, deconvolved TOM20 and TOM70 images were thresholded, and centroid-to-centroid distances were computed using the MorphoLibJ plugin in ImageJ.

### Cyto/Mito GFP accumulation analysis

Confocal image stacks were acquired with an Olympus FV3000 system equipped with a 60 × oil immersion objective, using sequential excitation to record DAPI, MTS-GFP, mCherry, HA (when present) and TOM20 channels. Raw files were imported into Fiji (Bio-Formats). Binary masks were generated by automated thresholding: the mCherry channel yielded the whole-cell mask, TOM20 the mitochondrial mask, and DAPI the nuclear mask. A cytosolic mask was produced by subtracting the nuclear and mitochondrial masks from the whole-cell mask (Image Calculator, “Subtract create”). Mean GFP fluorescence was then measured within (i) the cytosolic mask and (ii) the TOM20-defined mitochondrial mask. Import efficiency was expressed as the Cyto/Mito ratio:

Cyto/Mito ratio=mean GFP cytosol /mean GFP mitochondria

All threshold levels, arithmetic operations and measurement parameters were applied identically to every image and condition.

### Next-generation sequencing

Library preparation was done using the MGIEasy RNA library prep set (MGI) according to the manufacturer’s instructions. In brief, 500□ng total RNA was used as starting material from which rRNA was depleted using a Ribo-Zero Plus rRNA depletion kit (Illumina). The rRNA-depleted samples were fragmented and reverse transcribed, and the second strands were synthesized. DNA was then purified using DNA clean beads provided in the kit, followed by end repair and A-tailing. Barcoding and adaptor ligation were performed, and the samples were purified. Samples were amplified using adaptor-specific primers and quantified using a Qubit dsDNA high-sensitivity kit (Thermo Scientific). Sample fragment size was determined using a 4200 Tape Station (Agilent). The samples were denatured, and single-stranded circular DNA strands were generated. Further, rolling cycle amplification was performed to generate DNA nanoballs. The samples were subsequently loaded onto the flow cells (FCL) and paired end RNA sequencing was performed at MGI 2000 (MGI) platform for read length of 100 bases.

### Data processing and analysis

Raw sequencing reads were processed using Cutadapt(*64*) to remove MGI adapters and low-quality sequences. Reads with a quality score below 20 and a length shorter than 36 bp were discarded. The remaining high-quality reads were mapped to the human reference genome (GRCh38) using HISAT2 with default parameters(*65*). Uniquely aligned reads were quantified using the featureCounts function from the Subread package(*66*), based on a GTF annotation file downloaded from Ensembl(*67*), which contained read count data for 60,683 genes. Genes with fewer than 10 total read counts across all samples were excluded for further analysis. Differential gene expression analysis was performed using DESeq2(*68*), with genes considered differentially expressed if they had a P-value < 0.05 and |log2FC| ≥ 0.58.

### Functional enrichment analysis

Functional enrichment analysis was conducted using clusterProfiler(*69*) to identify enriched Gene Ontology (GO) terms. To refine the results, similar enriched terms were merged using the simplify function of ClusterProfiler, with a similarity cut-off of 0.5. The p-value was utilized as a criterion for selecting representative terms, while the min method was applied for feature selection. The Wang method was employed to measure similarity. Finally, the top five enriched GO terms for the genes were visualized.

### Statistical analysis

For each experiment, at least three independent replicates (until specified) were used to calculate mean ± SEM and plotted graphically wherever indicated. Statistical significance was measured using a two-tailed, unpaired Student t-test, and the resultant p values were represented as *, **, *** indicating p values ≤ 0.05, 0.01, and 0.001, respectively.

## Biosafety statement

All infection protocols in this study were approved by the Institute Biosafety Committee of CSIR-CCMB.

## Ethical Statement

All animal protocols in this study were approved by the Institute Animal Ethics Committee of CSIR-CCMB.

## Data availability

All data pertaining to this manuscript are included within the manuscript. The RNA sequencing (RNA-seq) data have been submitted to the Gene Expression Omnibus (GEO) database and are available under the accession number GSE293066.

## ACKNOWLEDGEMENTS

We thank Dr. Shasi Vardhan Kalivendi, Dr. Chitrakshi Pant and Abhijit Mohapatra for assistance with Seahorse Experiments. We also thank Dr. Nitesh Kumar Singh for help with the preliminary RNA-seq datasets. We thank Dr. Sudhanshu Vrati for sharing the mito-GFP-RFP plasmid. We thank Dr. Swasti Raychaudhuri for sharing the MTS-GFP plasmid. Special appreciation to Mohan Singh Moodu and Amit Kumar for their assistance with experimental logistics. We thank Doddetipalli Jaya Surya for his assistance with BN-PAGE experiments. We also thank Dr. C. Subbulakshmi, Dr. Venkata Mahesh Nitla and Suman Bandari for their help with confocal imaging and analysis, and N Sai Ram, Dr. M Jerald Mahesh Kumar and B Jyothi Lakshmi from CCMB Animal Facility.

## AUTHOR CONTRIBUTIONS

V.S and K.H.H conceived and designed this study. V.S, P.S.P, and K.H.H designed the experiments. V.S, P.S.P, K.S.N, and A.R.P performed the overexpression and KO studies. A.R.P and D.T. performed the IFN treatment studies. V.S and D.B generated MAVS KO Huh7 cells. P.P.S generated and verified the MAVS/IFNAR1 DKO mice. V.S, P.S.P, P.P.S, and D.T isolated MEFs for this study. V.S analysed the next-generation sequencing data with assistance from D.T. A.B.P, and K.S.V performed NMR experiments. S.C provided IFNAR1 KO mice. K.H.H and V.S wrote the manuscript with inputs from S.P, D.T, and P.P.S.

## Declaration of generative AI and AI-assisted technologies in the writing process

During the preparation of this work the author(s) used ChatGPT in order to improve language and readability. After using this tool/service, the author(s) reviewed and edited the content as needed and take(s) full responsibility for the content of the publication.

## REFERENCES

1. N. Yan, Z. J. Chen, Intrinsic antiviral immunity. Nature Immunology 13, 214–222 (2012).

2. P. D. Bieniasz, Intrinsic immunity: a front-line defense against viral attack. Nature Immunology 5, 1109–1115 (2004).

3. J. Moretti, J. M. Blander, Cell-autonomous stress responses in innate immunity. Journal of Leukocyte Biology 101, 77–86 (2017).

4. S. Majdoul, A. A. Compton, Lessons in self-defence: inhibition of virus entry by intrinsic immunity. Nature Reviews Immunology 22, 339–352 (2022).

5. N. K. Duggal, M. Emerman, Evolutionary conflicts between viruses and restriction factors shape immunity. Nature Reviews Immunology 12, 687–695 (2012).

6. R. B. Seth, L. Sun, C. K. Ea, Z. J. Chen, Identification and characterization of MAVS, a mitochondrial antiviral signaling protein that activates NF-κB and IRF3. Cell 122, 669–682 (2005).

7. E. Dixit et al., Peroxisomes Are Signaling Platforms for Antiviral Innate Immunity. Cell 141, 668–681 (2010).

8. F. Hou et al., MAVS forms functional prion-like aggregates to activate and propagate antiviral innate immune response. Cell 146, 448–461 (2011).

9. S. M. Belgnaoui, S. Paz, J. Hiscott, Orchestrating the interferon antiviral response through the mitochondrial antiviral signaling (MAVS) adapter. Current Opinion in Immunology 23, 564–572 (2011).

10. D. Tandel et al., SARS-CoV-2 Variant Delta Potently Suppresses Innate Immune Response and Evades Interferon-Activated Antiviral Responses in Human Colon Epithelial Cells. Microbiology Spectrum 10, e01604–01622 (2022).

11. C.-S. Shi et al., SARS-Coronavirus Open Reading Frame-9b Suppresses Innate Immunity by Targeting Mitochondria and the MAVS/TRAF3/TRAF6 Signalosome. The Journal of Immunology 193, 3080–3089 (2014).

12. E. Meylan, Cardif is an adaptor protein in the RIG-I antiviral pathway and is targeted by hepatitis C virus. Nature 437, 1167–1172 (2005).

13. C. Li et al., E3 ubiquitin ligase MARCH5 positively regulates Japanese encephalitis virus infection by catalyzing the K27-linked polyubiquitination of viral E protein and inhibiting MAVS-mediated type I interferon production. mBio 16, e00208–00225 (2025).

14. Z. He et al., Dengue Virus Subverts Host Innate Immunity by Targeting Adaptor Protein MAVS. Journal of Virology 90, 7219–7230 (2016).

15. A. P. West, G. S. Shadel, S. Ghosh, Mitochondria in innate immune responses. Nature Reviews Immunology 11, 389–402 (2011).

16. E. Marques, R. Kramer, D. G. Ryan, Multifaceted mitochondria in innate immunity. npj Metabolic Health and Disease 2, 6 (2024).

17. C. Shang et al., SARS-CoV-2 Causes Mitochondrial Dysfunction and Mitophagy Impairment. Frontiers in Microbiology Volume 12 **-** 2021, (2022).

18. K. Yang et al., Japanese encephalitis virus infection induces mitochondrial-mediated apoptosis through the proapoptotic protein BAX. Frontiers in Microbiology Volume 15 **-** 2024, (2024).

19. L. Chatel-Chaix et al., Dengue Virus Perturbs Mitochondrial Morphodynamics to Dampen Innate Immune Responses. Cell Host & Microbe 20, 342–356 (2016).

20. M. Khan, G. H. Syed, S.-J. Kim, A. Siddiqui, Mitochondrial dynamics and viral infections: A close nexus. Biochimica et Biophysica Acta (BBA) - Molecular Cell Research 1853, 2822–2833 (2015).

21. M. Sorouri, T. Chang, C. Hancks Dustin, Mitochondria and Viral Infection: Advances and Emerging Battlefronts. mBio 13, e02096–02021 (2022).

22. A. Agarwal et al., Japanese Encephalitis Virus NS4A Protein Interacts with PTEN-Induced Kinase 1 (PINK1) and Promotes Mitophagy in Infected Cells. Microbiology Spectrum 10, e00830–00822 (2022).

23. V. Barbier, D. Lang, S. Valois, A. L. Rothman, Carey L. Medin, Dengue virus induces mitochondrial elongation through impairment of Drp1-triggered mitochondrial fission. Virology 500, 149–160 (2017).

24. B. Singh et al., Defective Mitochondrial Quality Control during Dengue Infection Contributes to Disease Pathogenesis. Journal of Virology 96, e00828–00822 (2022).

25. S.-J. Kim et al., Hepatitis C virus triggers mitochondrial fission and attenuates apoptosis to promote viral persistence. Proceedings of the National Academy of Sciences 111, 6413–6418 (2014).

26. S. Lenhard et al., The Orf9b protein of SARS-CoV-2 modulates mitochondrial protein biogenesis. Journal of Cell Biology 222, e202303002 (2023).

27. S. M. Horner, H. M. Liu, H. S. Park, J. Briley, M. Gale, Mitochondrial-associated endoplasmic reticulum membranes (MAM) form innate immune synapses and are targeted by hepatitis C virus. Proceedings of the National Academy of Sciences of the United States of America 108, 14590–14595 (2011).

28. T. Hayashi, R. Rizzuto, G. Hajnoczky, T. P. Su, MAM: more than just a housekeeper. Trends in Cell Biology 19, 81–88 (2009).

29. X.-Y. Liu, B. Wei, H.-X. Shi, Y.-F. Shan, C. Wang, Tom70 mediates activation of interferon regulatory factor 3 on mitochondria. Cell Research 20, 994–1011 (2010).

30. X. Gao et al., Crystal structure of SARS-CoV-2 Orf9b in complex with human TOM70 suggests unusual virus-host interactions. Nature Communications 12, 2843 (2021).

31. R. Lin, S. Paz, J. Hiscott, Tom70 imports antiviral immunity to the mitochondria. Cell Research 20, 971–973 (2010).

32. L. D. Zorova et al., Mitochondrial membrane potential. Analytical Biochemistry 552, 50–59 (2018).

33. A. C. Y. Fan et al., Interaction between the Human Mitochondrial Import Receptors Tom20 and Tom70 *in Vitro* Suggests a Chaperone Displacement Mechanism * . Journal of Biological Chemistry 286, 32208–32219 (2011).

34. X.-D. Li, L. Sun, R. B. Seth, G. Pineda, Z. J. Chen, Hepatitis C virus protease NS3/4A cleaves mitochondrial antiviral signaling protein off the mitochondria to evade innate immunity. Proceedings of the National Academy of Sciences 102, 17717–17722 (2005).

35. W. Sun et al., Antiviral Adaptor MAVS Promotes Murine Lupus With a B Cell Autonomous Role. Frontiers in Immunology Volume 10 - 2019, (2019).

36. Y.-T. Kao, M. M. C. Lai, C.-Y. Yu, How Dengue Virus Circumvents Innate Immunity. Frontiers in Immunology Volume 9 **-** 2018, (2018).

37. Y.-G. Zhang et al., Type I/type III IFN and related factors regulate JEV infection and BBB endothelial integrity. Journal of Neuroinflammation 20, 216 (2023).

38. D. Vedagiri et al., Retinoic Acid-Inducible Gene I-Like Receptors Activate Snail To Limit RNA Viral Infections. Journal of Virology 95, 10.1128/jvi.01216-01221 (2021).

39. J. Deng et al., SARS-CoV-2 NSP7 inhibits type I and III IFN production by targeting the RIG-I/MDA5, TRIF, and STING signaling pathways. Journal of Medical Virology 95, e28561 (2023).

40. R. S. Freitas, T. F. Crum, K. Parvatiyar, SARS-CoV-2 Spike Antagonizes Innate Antiviral Immunity by Targeting Interferon Regulatory Factor 3. Frontiers in Cellular and Infection Microbiology Volume 11 - 2021, (2022).

41. J. M. Minkoff, B. tenOever, Innate immune evasion strategies of SARS-CoV-2. Nature Reviews Microbiology 21, 178–194 (2023).

42. Y. Hanada et al., MAVS is energized by Mff which senses mitochondrial metabolism via AMPK for acute antiviral immunity. Nature Communications 11, 5711 (2020).

43. R. Jin et al., Japanese Encephalitis Virus Activates Autophagy as a Viral Immune Evasion Strategy. PLOS ONE 8, e52909 (2013).

44. D. Zhou et al., The Japanese Encephalitis Virus NS1′ Protein Inhibits Type I IFN Production by Targeting MAVS. The Journal of Immunology 204, 1287–1298 (2020).

45. D. Zhou et al., The Japanese Encephalitis Virus NS1’ Protein Inhibits Type I IFN Production by Targeting MAVS.

46. A. Dalrymple Nadine, V. Cimica, R. Mackow Erich, Dengue Virus NS Proteins Inhibit RIG-I/MAVS Signaling by Blocking TBK1/IRF3 Phosphorylation: Dengue Virus Serotype 1 NS4A Is a Unique Interferon-Regulating Virulence Determinant. mBio 6, 10.1128/mbio.00553-00515 (2015).

47. A. A.-O. Agarwal et al., Japanese Encephalitis Virus NS4A Protein Interacts with PTEN-Induced Kinase 1 (PINK1) and Promotes Mitophagy in Infected Cells.

48. F. Le Chevalier et al., Mice humanized for MHC and hACE2 with high permissiveness to SARS-CoV-2 omicron replication. Microbes and Infection 25, 105142 (2023).

49. H. Chu et al., Comparative tropism, replication kinetics, and cell damage profiling of SARS-CoV-2 and SARS-CoV with implications for clinical manifestations, transmissibility, and laboratory studies of COVID-19: an observational study. The Lancet Microbe 1, e14–e23 (2020).

50. W.-H. Shao et al., Prion-like Aggregation of Mitochondrial Antiviral Signaling Protein in Lupus Patients Is Associated With Increased Levels of Type I Interferon. Arthritis & Rheumatology 68, 2697–2707 (2016).

51. S. Biacchesi et al., Mitochondrial Antiviral Signaling Protein Plays a Major Role in Induction of the Fish Innate Immune Response against RNA and DNA Viruses. Journal of Virology 83, 7815–7827 (2009).

52. C. Kwak et al., Contact-ID, a tool for profiling organelle contact sites, reveals regulatory proteins of mitochondrial-associated membrane formation. Proceedings of the National Academy of Sciences 117, 12109–12120 (2020).

53. S. M. Horner, H. M. Liu, H. S. Park, J. Briley, M. Gale, Mitochondrial-associated endoplasmic reticulum membranes (MAM) form innate immune synapses and are targeted by hepatitis C virus. Proceedings of the National Academy of Sciences 108, 14590–14595 (2011).

54. J. L. Jacobs, J. Zhu, S. N. Sarkar, C. B. Coyne, Regulation of Mitochondrial Antiviral Signaling (MAVS) Expression and Signaling by the Mitochondria-associated Endoplasmic Reticulum Membrane (MAM) Protein Gp78 *. Journal of Biological Chemistry 289, 1604–1616 (2014).

55. N. S. Gokhale et al., Cellular RNA interacts with MAVS to promote antiviral signaling. Science 386, eadl0429.

56. S.-J. Kim et al., Hepatitis B Virus Disrupts Mitochondrial Dynamics: Induces Fission and Mitophagy to Attenuate Apoptosis. PLOS Pathogens 9, e1003722 (2013).

57. S. Rawat, V. Anusha, M. Jha, K. Sreedurgalakshmi, S. Raychaudhuri, Aggregation of Respiratory Complex Subunits Marks the Onset of Proteotoxicity in Proteasome Inhibited Cells. Journal of Molecular Biology 431, 996–1015 (2019).

58. R. Gosert et al., Identification of the Hepatitis C Virus RNA Replication Complex in Huh-7 Cells Harboring Subgenomic Replicons. Journal of Virology 77, 5487–5492 (2003).

59. J. Brugarolas, R. T. Bronson, T. Jacks, p21 Is a Critical CDK2 Regulator Essential for Proliferation Control in Rb-deficient Cells. Journal of Cell Biology 141, 503–514 (1998).

60. P. Gupta et al., Kingdom-specific lipid unsaturation calibrates sequence evolution in membrane arm subunits of eukaryotic respiratory complexes. Nature Communications 16, 2044 (2025).

61. A. B. Patel, D. L. Rothman, G. W. Cline, K. L. Behar, Glutamine is the major precursor for GABA synthesis in rat neocortex in vivo following acute GABA-transaminase inhibition. Brain Research 919, 207–220 (2001).

62. P. Bagga, A. N. Chugani, K. S. Varadarajan, A. B. Patel, In vivo NMR studies of regional cerebral energetics in MPTP model of Parkinson’s disease: recovery of cerebral metabolism with acute levodopa treatment. Journal of Neurochemistry 127, 365–377 (2013).

63. P. K. Mishra et al., Impaired neuronal and astroglial metabolic activity in chronic unpredictable mild stress model of depression: Reversal of behavioral and metabolic deficit with lanicemine. Neurochemistry International 137, 104750 (2020).

64. M. Martin, Cutadapt removes adapter sequences from high-throughput sequencing reads. EMBnet.journal; Vol 17, No 1: Next Generation Sequencing Data Analysis, (2011).

65. D. Kim, J. M. Paggi, C. Park, C. Bennett, S. L. Salzberg, Graph-based genome alignment and genotyping with HISAT2 and HISAT-genotype. Nature Biotechnology 37, 907–915 (2019).

66. Y. Liao, G. K. Smyth, W. Shi, featureCounts: an efficient general purpose program for assigning sequence reads to genomic features. Bioinformatics 30, 923–930 (2014).

67. K. L. Howe et al., Ensembl 2021. Nucleic Acids Research 49, D884–D891 (2021).

68. M. I. Love, W. Huber, S. Anders, Moderated estimation of fold change and dispersion for RNA-seq data with DESeq2. Genome Biology 15, 550 (2014).

69. T. Wu et al., clusterProfiler 4.0: A universal enrichment tool for interpreting omics data. The Innovation 2, 100141 (2021).

