## Supplementary figures for "MAVS Safeguards Mitochondrial Integrity to Drive a Potent Intrinsic Antiviral Program"

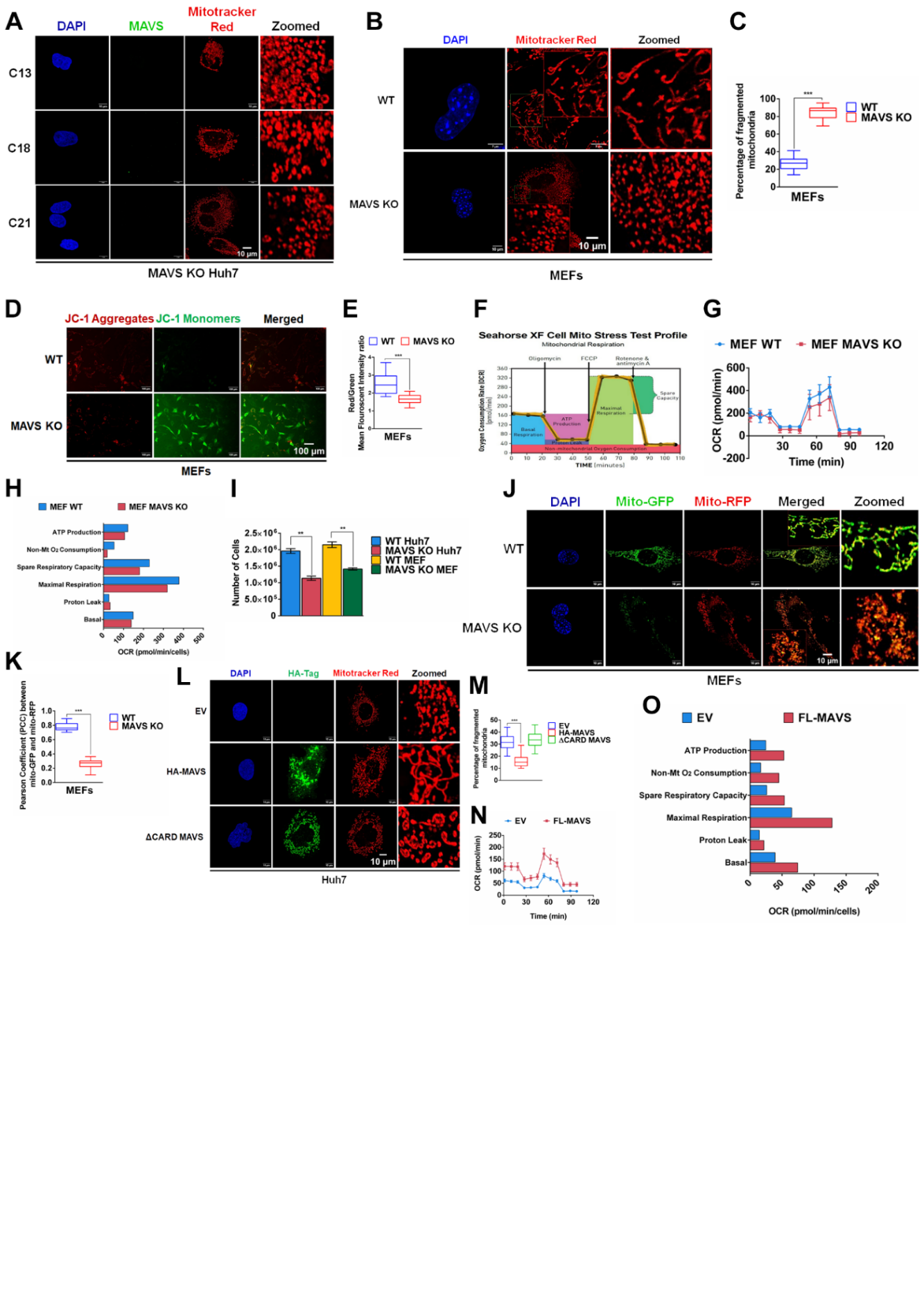

**Figure S1. MAVS is critical for maintaining mitochondrial structure and homeostasis**

**(A)** Confocal microscopy images of mitochondrial fragmentation in three different clones of MAVS KO Huh7. MAVS was knocked out of Huh7 cells using the CRISPR-Cas9 method. Three colonies (C13, C18, and C21) were screened for phenotype confirmation. C13 was used for the subsequent studies. **(B)** Confocal microscopy images of mitochondrial fragmentation in WT and MAVS KO MEFs, and **(C)** quantitative representation of mitochondrial fragmentation from images in (B). **(D)** Epifluorescence microscopy images analyzing mitochondrial membrane potential in WT and MAVS KO MEFs. Scale bars = 100 µm, and **(E)** Quantitative representation of mitochondrial membrane potential from images in (D) (n = 100 cells). (**F**) Schematic of a mitochondrial stress assay (**G**) Graphs showing OCR and **(H)** mitochondrial respiration parameters in WT and MAVS KO MEFs. (**I**) Comparison of cell division in WT and MAVS KO Huh7 cells and MEFs. Equal number of cells were initially plated and incubated for 48 hours. Following this, the cells were trypsinized and counted using a hemocytometer. **(J)** Confocal microscopy images analyzing mitophagy in WT and MAVS KO MEFs, and **(K)** the quantitative representation of mitophagy from images in (J). **(L)** Confocal microscopy images of mitochondrial fragmentation in uninfected Huh7 cells transfected with FL-MAVS or ΔCARD MAVS, and **(M)** quantitative representation of mitochondrial fragmentation from images in (L). (**N**) Graphs showing OCR and **(O)** mitochondrial respiration parameters in uninfected Huh7 cells transfected with FL-MAVS. Scale bars, 10 µm and n = 50 cells unless otherwise indicated. Data are presented as mean ± SEM, with *, ** and *** denoting p <0.05, 0.01 and < 0.001.

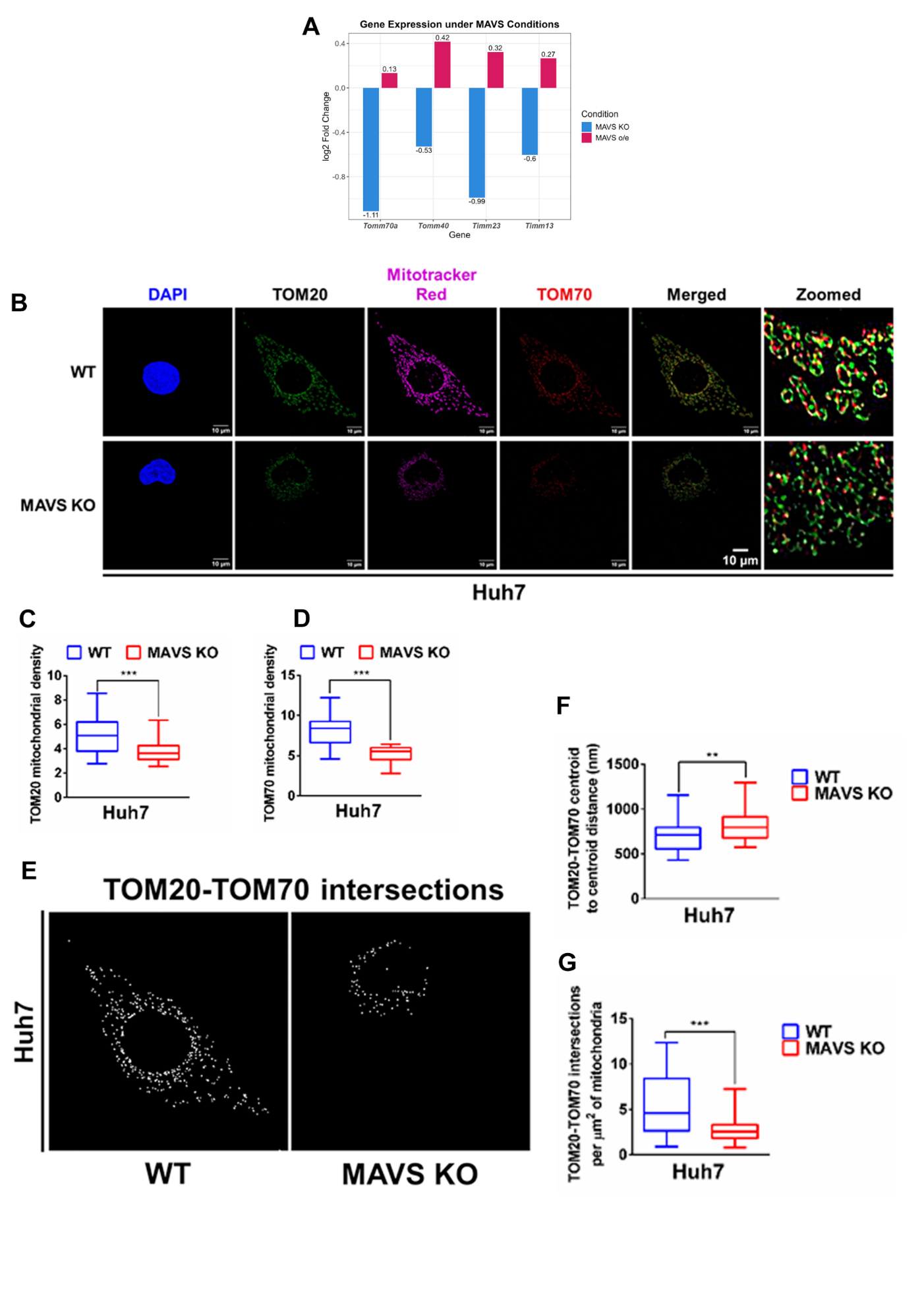

**Figure S2. MAVS safeguards mitochondria by stabilizing TOM complex and protein import**

**(A)** Bar graph showing DEGs associated with TIM and TOM complex under MAVS overexpressed and MAVS deficient conditions. **(B)** Confocal images showing co-localization of TOM20 and TOM70 puncta in WT and MAVS KO Huh7 cells. **(C)** Graph showing mitochondrial density of TOM20, and **(D)** TOM70. **(E)** Intersection masks highlighting TOM20 and TOM70 overlaps, and **(F)** quantification of intersection density per unit mitochondrial area. **(G)** Quantification of centroid-to-centroid distances between TOM20 and TOM70 in WT and MAVS KO cells. Scale bars, 10 µm and n = 50 cells unless otherwise indicated. Data are presented as mean ± SEM, with *, ** and *** denoting p <0.05, 0.01 and < 0.001.

**Figure S3**

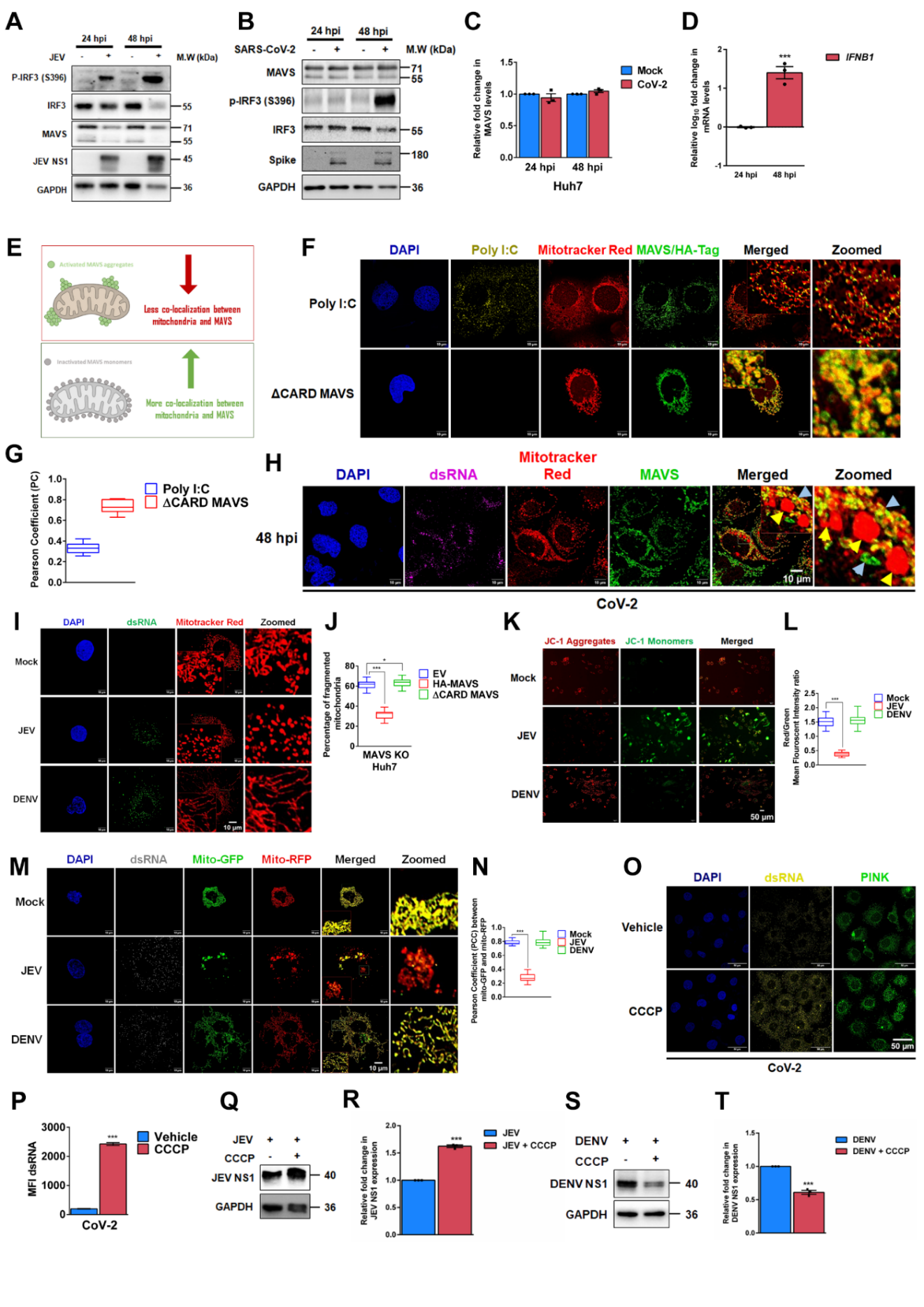

**Figure S3. Viruses target MAVS aggregation to destabilize mitochondrial homeostasis**

**(A)** Immunoblot analysis of IRF-3 phosphorylation, MAVS, and JEV NS1 levels in Huh7 cells infected with JEV at different time points. **(B)** Immunoblot analyzing MAVS, IRF-3 phosphorylation, and SARS-CoV-2 Spike in Huh7 cells infected with SARS-CoV-2 at different time points. **(C)** Densitometric analysis of MAVS expression levels in Huh7 cells upon CoV-2 infection at different time points. (**D**) qRT-PCR analysis showing relative fold change in *IFNB1* mRNA levels in Huh7 cells at 24 and 48 hpi. (**E**) Schematic depicting MAVS aggregation and its foci formation on mitochondria. (**F**) Confocal microscopy images showing changes in MAVS aggregation upon transection with Poly (I: C) or aggregation-deficient HA-tagged ΔCARD MAVS, stained with Mitotracker Red, fixed, and subjected to endogenous staining for HA and MAVS. **(G)** Quantification of the PC for MAVS and mitochondria co-localization from images in (F) (n = 250 ROIs). **(H)** Confocal microscopy panel showing mitochondrial fragmentation in Huh7 cells infected with SARS-CoV-2 at 48 hpi. Yellow arrow in zoomed inset image shows MAVS deficient mitochondria, whereas blue arrows shows mitochondria with intact MAVS **(I)** Confocal microscopy images analyzing mitochondrial fragmentation in Huh7 cells infected with JEV and DENV using Mitotracker Red, and **(J)** quantification of mitochondrial fragmentation from images in (I). **(K)** Epifluorescence microscopy images showing mitochondrial depolarization in Huh7 cells infected with JEV and DENV using JC1 dye (Scale bars = 50 µm) and (**L**) quantitative representation of mitochondrial membrane potential from images in (K) (n = 100 cells). **(M)** Confocal microscopy images analyzing mitophagy using mito-GFP-RFP in Huh7 cells infected with JEV and DENV, and **(N)** quantitative representation of mitophagy from images in (M). **(O)** Confocal microscopy images detecting dsRNA and PINK1 in CoV-2 infected cells upon treatment with CCCP and **(P)** Mean Fluorescent Intensity (MFI) quantification of dsRNA signal intensity in these cells (n = 100 cells). **(Q)** Immunoblot analysis showing JEV NS1 expression in CCCP-treated Huh7 cells and (**R**) Densitometric analysis of JEV NS1 expression in (Q). **(S)** Immunoblot analysis showing DENV NS1 expression in CCCP-treated Huh7 cells and (**T**) Densitometric analysis of DENV NS1 expression in (S). Scale bars = 10 µm and n = 50 cells unless mentioned otherwise. Data are presented as mean ± SEM, with *, ** and *** denoting p <0.05, 0.01 and < 0.001.

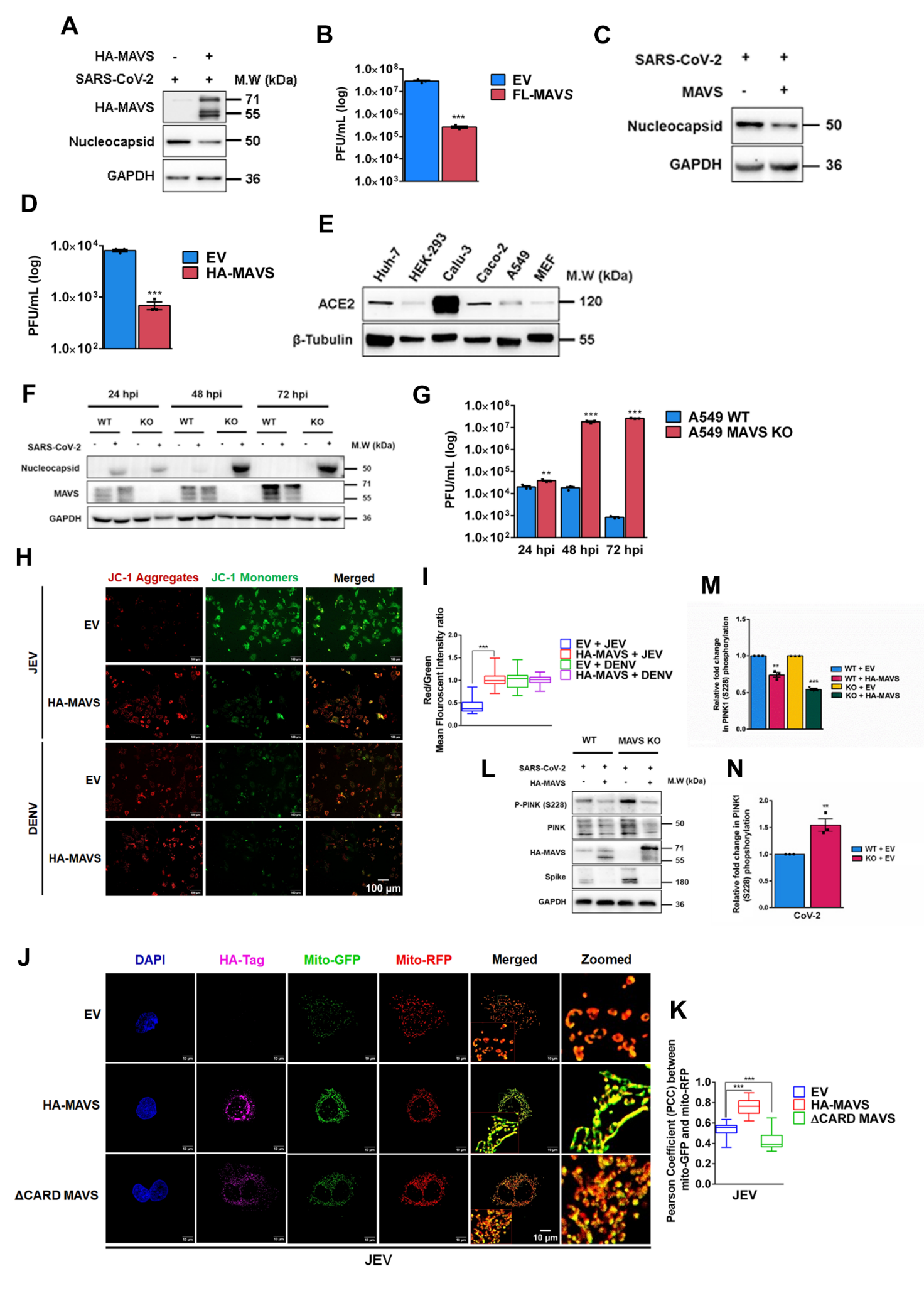

**Figure S4. MAVS implements mitochondrial homeostasis to restrict RNA virus replication**

**(A)** Immunoblot analysis of SARS-CoV-2 nucleocapsid protein expression upon overexpression of FL-MAVS in Caco2 cells, and (**B**) plaque assay quantification of infectious viral titers from their supernatants. **(C)** Immunoblot analysis of SARS-CoV-2 nucleocapsid protein expression upon overexpression of full-length MAVS (FL-MAVS) in A549 MAVS KO cells infected with SARS-CoV-2, and (**D**) plaque assay quantification of infectious viral titers from their supernatants. **(E)** Immunoblot showing ACE2 levels across different cell types. **(F)** Immunoblot analysing SARS-CoV-2 nucleocapsid levels in WT and MAVS KO MEFs infected with SARS-CoV-2. **(G)** Infectious viral titers comparing infection in WT and MAVS KO MEFs. **(H)** Epifluorescence microscopy images analyzing mitochondrial membrane potential in JEV or DENV-infected Huh7 cells upon supplementation of FL-MAVS. Scale bars = 100 µm, and **(I)** Quantitative representation of mitochondrial membrane potential from images in (H) (n = 100 cells). **(J)** Confocal microscopy images analyzing mitophagy using mito-GFP-RFP in Huh7 cells supplemented with HA-tagged FL-MAVS or ΔCARD-MAVS upon infection with JEV, and **(K)** the quantitative representation of mitophagy from images in (J). **(L)** Immunoblot analysis of PINK1 phosphorylation, HA-MAVS, and SARS-CoV-2 Spike in WT and MAVS KO A549 cells expressing FL-MAVS. (**M**) Quantitative densitometric analysis showing relative fold change in PINK1 phosphorylation between empty vector and HA-MAVS transfected cells, and (**N**) comparing the empty vector-transfected WT and MAVS KO A549 cells. Scale bars = 10 µm and n = 50 cells unless mentioned otherwise. Data are presented as mean ± SEM, with *, ** and *** denoting p <0.05, 0.01 and < 0.001.

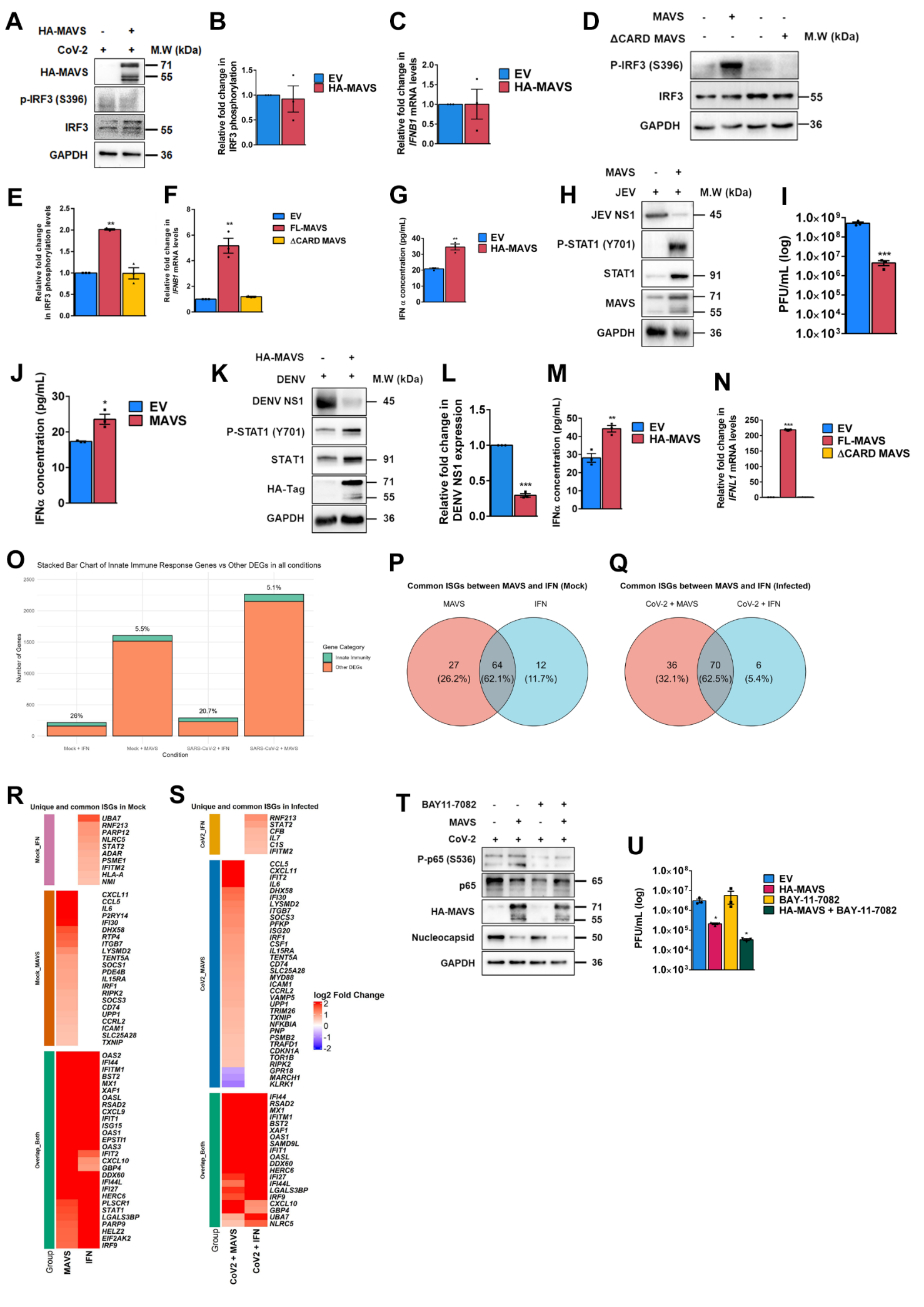

**Figure S5. MAVS restricts viral infection through an interferon-independent intrinsic antiviral program**

(**A**) Immunoblot analysis of IRF-3 phosphorylation in Caco2 cells supplemented with FL-MAVS, followed by SARS-CoV-2 infection. (**B**) Densitometric quantification of IRF-3 phosphorylation from (A). (**C**) qRT-PCR analysis showing relative fold change in *IFNB1* mRNA levels in Caco2 cells supplemented with FL-MAVS, followed by SARS-CoV-2 infection. (**D**) Immunoblot analysis of IRF-3 phosphorylation in uninfected Huh7 cells supplemented with FL-MAVS and ΔCARD MAVS. (**E**) Densitometric quantification of IRF3 phosphorylation from (D). (**F**) qRT-PCR analysis showing relative fold change in *IFNB1* mRNA levels in uninfected Huh7 cells supplemented with FL-MAVS and ΔCARD MAVS. (**G**) Relative fold change in IFN-α concentrations secreted by uninfected Huh7 cells supplemented with FL-MAVS. (**H**) Immunoblot analysis of JEV NS1 expression, STAT1 phosphorylation, and MAVS expression in Huh7 cells supplemented with FL-MAVS, followed by JEV expression and (**I**) plaque assay quantification of infectious viral titers from their supernatants. (**J**) Relative fold change in IFN-α concentrations in Huh7 supernatant supplemented with FL-MAVS followed by JEV infection. (**K**) Immunoblot analyzing DENV NS1 expression, STAT1 phosphorylation, and HA-MAVS expression in Huh7 cells supplemented with FL-MAVS, followed by DENV infection and (**L**) Densitometric quantification of DENV NS1 expression from (K). (**M**) Relative fold change in IFN-α concentrations in Huh7 supernatant supplemented with FL-MAVS followed by DENV infection. (**N**) qRT-PCR analysis showing relative fold change in *IFNL1* mRNA levels in uninfected Huh7 cells supplemented with FL-MAVS and ΔCARD MAVS. **(O)** Stacked bar chart showing the percentage of DEGs associated with innate immunity in all conditions involving MAVS overexpression and IFN-α treatment. The green stack indicates genes associated with innate immunity, and the orange stack denotes other DEGs. **(P)** Venn diagram showing overlap of ISGs between MAVS and IFN-α treated samples in uninfected and **(Q)** SARS-CoV-2 infected conditions. **(R)** Heatmap showing expression levels of overlapping ISGs and exclusive ISGs to MAVS and IFN-α treatment in uninfected and **(S)** CoV-2-infected conditions. Expression values are represented as Log_2_ Fold Change. **(T)** Immunoblot showing p65 phosphorylation, MAVS, and nucleocapsid levels in SARS-CoV-2 infected Huh7 cells overexpressing MAVS or treated with BAY11-7082, and **(U)** infectious viral titers. Data are presented as mean ± SEM, with *, ** and *** denoting p <0.05, 0.01 and < 0.001.

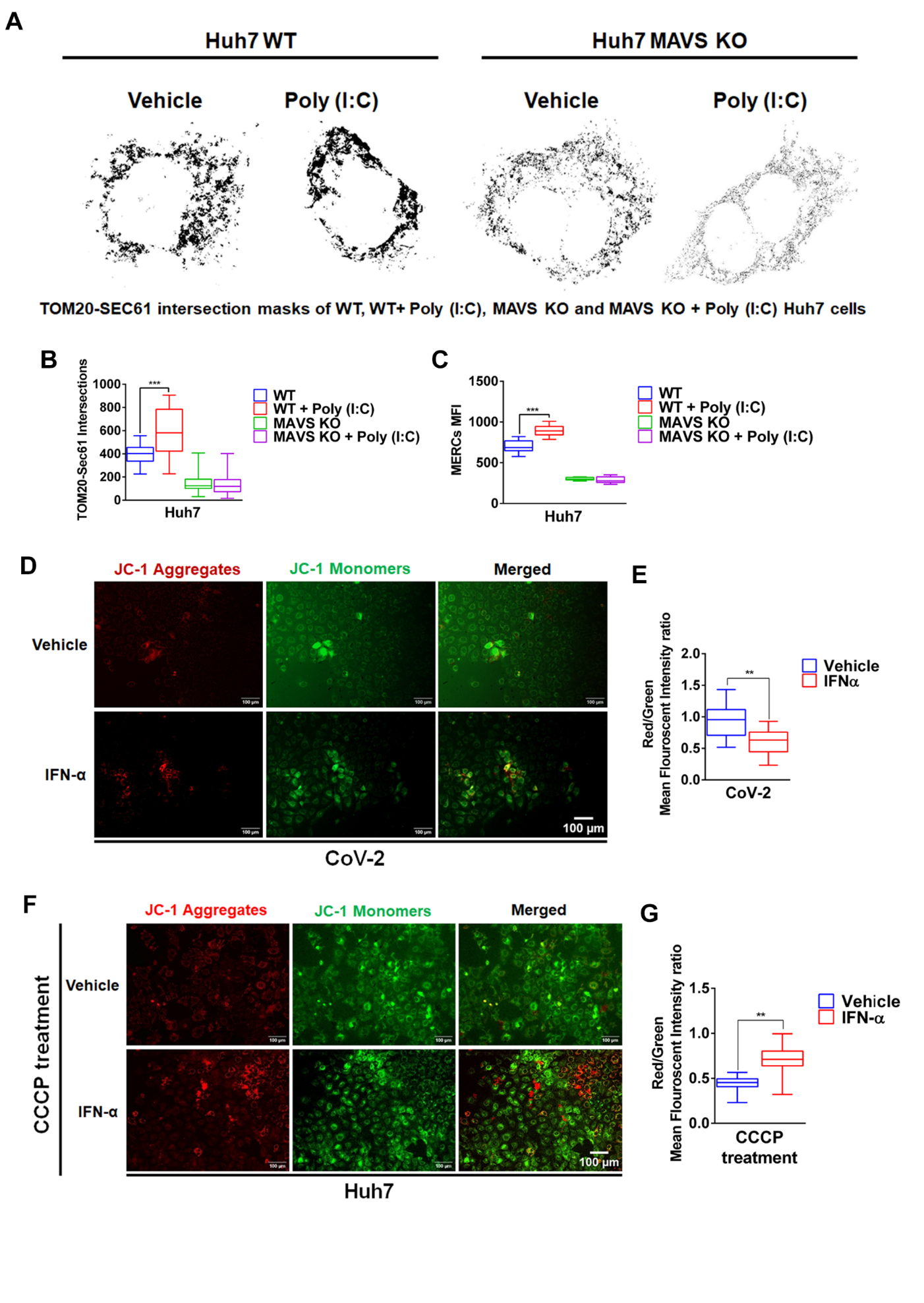

**Figure S6. MAVS-mediated mitochondrial safeguarding is not contingent upon IFN signaling**

**(A)** Intersection masks highlighting HA-TOM20 (mitochondria) and FLAG-Sec61 (ER) overlaps, **(B)** quantification of intersection masks and **(C)** MAM intensity in the vehicle or Poly(I:C) treated WT and MAVS KO cells. **(D)** Epifluorescence microscopy images analyzing mitochondrial membrane potential in SARS-CoV-2 infected Huh7 cells upon IFN-α treatment, and **(E)** Quantitative representation of mitochondrial membrane potential from images in (D) (n = 100 cells, Scale 100 µm). **(F)** Epifluorescence microscopy images analyzing mitochondrial membrane potential in CCCP-treated Huh7 cells upon IFN-α treatment, and **(G)** Quantitative representation of mitochondrial membrane potential from images in (F) (n = 100 cells, Scale 100 µm). Scale bars = 10 µm and n = 50 cells unless mentioned otherwise. Data are presented as mean ± SEM, with *, ** and *** denoting p <0.05, 0.01 and < 0.001.
